# SWI/SNF nuclear foci scaffold peripheral gene hubs for circadian chromatin control

**DOI:** 10.64898/2026.02.03.703661

**Authors:** Qianqian Chen, Ye Yuan, Dunham Clark, Swathi Yadlapalli

**Affiliations:** Department of Cell and Developmental Biology, University of Michigan, Ann Arbor, MI 48109; Cellular and Molecular Biology Graduate Program, University of Michigan, Ann Arbor, MI 48109

## Abstract

How dynamic circadian rhythms arise from a relatively static nuclear architecture remains a fundamental open question. Here, we identify the SWI/SNF chromatin-remodeling complex as a critical interface between the circadian clock and nuclear organization. Using endogenously tagged Moira (MOR), a core component of the Drosophila SWI/SNF complex and homolog of human BAF155, we find that MOR assembles into a small number of discrete foci near the nuclear periphery, in stark contrast to the current model of diffuse nuclear distribution for SWI/SNF-family proteins. We demonstrate that this localization is maintained by the inner nuclear envelope LEM-domain protein Otefin, effectively shifting MOR from a global to a localized regulator of chromatin architecture. DNA-FISH reveals that clock-regulated genes cluster into peripheral “hubs” that co-localize with MOR foci throughout the circadian cycle. ATAC-seq analysis shows that while MOR modulates chromatin accessibility genome-wide, it establishes a constitutive restrictive baseline specifically at clock-regulated loci. As a result, MOR depletion abolishes accessibility rhythms at these loci, rendering them constitutively hyper-accessible. This deregulation disrupts rhythmic gene expression and ultimately drives behavioral arrhythmia. Strikingly, oscillations of the core clock proteins PER and TIM remain intact, indicating that MOR loss uncouples the protein oscillator from its genomic output. Together, these results reveal that MOR-containing SWI/SNF foci form a stable perinuclear scaffold that gates chromatin accessibility, enabling the core clock machinery to convert transient protein oscillations into high-amplitude transcriptional rhythms.

**Significance Statement:** How dynamic circadian rhythms emerge from a relatively static nuclear architecture remains a fundamental mystery. Here, we challenge the assumption that clock-target genes are randomly distributed by identifying a precise nuclear organization where these genes cluster into peripheral hubs. These hubs co-localize with stable SWI/SNF (MOR) foci anchored at the nuclear envelope. We demonstrate that this perinuclear scaffold is essential for gating rhythmic chromatin accessibility. Most importantly, loss of this architecture uncouples robustly oscillating clock protein program from its genomic and behavioral outputs. These findings reveal that circadian timekeeping is not merely a biochemical process but a spatially organized one, where nuclear topology licenses the translation of time into circadian rhythms.

## Introduction

Circadian clocks are cell-autonomous molecular oscillators that synchronize physiology and behavior with daily environmental cycles^1^. In animals, these clocks are built upon transcriptional-translational feedback loops (TTFLs) in which the transcriptional activators CLOCK (CLK) and CYCLE (CYC) drive rhythmic expression of core clock genes such as *period* (*per*) and *timeless* (*tim*). The proteins encoded by these genes, PER and TIM, accumulate, dimerize, and feed back to repress CLK/CYC activity, thereby closing the feedback loop with ∼24-hour periodicity. Through this rhythmic regulation, the circadian system controls the expression of hundreds of downstream targets that coordinate neuronal activity, metabolism, and behavior^1, 2^.

While the temporal logic of this transcriptional program is well characterized, it remains unclear how these dynamics interface with the three-dimensional (3D) organization of the genome. Global nuclear architecture, including chromatin compartments and lamina-associated domains (LADs), is generally considered to be stable over time^3^. This global stability stands in contrast to the local, high-magnitude oscillations in histone modifications^4, 5^, chromatin accessibility^6, 7^, and enhancer-promoter interactions^8^ observed over the circadian cycle. It has long been a mystery how clock-regulated genes are embedded within and constrained by this relatively static architectural landscape.

This question is made particularly compelling by our recent finding that many clock-regulated genes reside at the nuclear periphery throughout the circadian cycle^9, 10^. While the periphery is traditionally viewed as a zone of constitutive silencing^11^, clock genes located at the periphery must remain poised for rhythmic activation and repression, enabling high-amplitude oscillations in gene expression. Despite emerging evidence for spatial compartmentalization among clock components^9, 12^, the mechanisms by which these spatial and temporal dimensions integrate to generate robust circadian timing remain incompletely understood.

To identify factors that might couple the core clock components to chromatin and nuclear architecture, we recently performed BioID proximity labeling^13^ using PERIOD (PER) as bait, reasoning that as the primary nuclear repressor, PER recruits the machinery required to enforce rhythmic silencing. We found that PER associates with the SWI/SNF chromatin-remodeling complex specifically during the repression phase of the circadian cycle (manuscript in preparation). SWI/SNF complexes, which are ATP-dependent chromatin remodelers, are structurally heterogeneous and functionally versatile^14–16^: while classically viewed as chromatin openers that mobilize nucleosomes to activate gene expression^17, 18^, they can also enforce repression by positioning nucleosomes to block transcription start sites^19–22^. Consistent with this functional diversity, SWI/SNF complexes are mutated in >20% of human cancers^14^

This functional versatility of SWI/SNF complexes extends to circadian regulation^23–25^. In *Neurospora crassa*, the binding of the WCC to C-box sequences near the *frq* promoter triggers dynamic nucleosome remodeling; specifically, the recruitment of SWI/SNF remodelers by WC-1 leads to nucleosome eviction, driving transcriptional activation of the *frq* gene^23^. Conversely, in *Drosophila*, the SWI/SNF complex has been shown to inhibit CLK/CYC-mediated transcription by promoting a repressive chromatin state and increasing nucleosome occupancy at core clock promoters^24^. However, whether SWI/SNF complexes function beyond local chromatin remodeling to act as higher-order spatial or architectural “gates” for circadian programs remains unexplored.

Here, we investigate the role of nuclear architecture in circadian control by focusing on Moira (MOR), a core structural subunit of the *Drosophila* SWI/SNF complex^26–28^. We reveal that MOR forms discrete foci at the nuclear periphery in *Drosophila* clock neurons, anchored by the inner nuclear envelope protein Otefin^29^, a LEM domain protein. Clock-regulated genes cluster within these MOR foci throughout the circadian cycle. Functionally, MOR restricts chromatin accessibility at these loci; its depletion results in constitutively open chromatin and loss of rhythmic gene expression. Consequently, MOR loss causes behavioral arrhythmicity despite intact PER/TIM oscillations, effectively uncoupling the molecular clock from its output. Together, our results reveal that clock genes are organized into SWI/SNF-dependent perinuclear hubs, providing an architectural scaffold that enables chromatin accessibility rhythms and drives circadian output.

## Results

### SWI/SNF subunit Moira is organized into discrete nuclear foci at the nuclear periphery

Previous work from our group has shown that chromatin accessibility in *Drosophila* clock neurons is under robust circadian control, exhibiting significant oscillations at the promoters and enhancers of hundreds of genes^6^. Given recent evidence from our BioID experiments that PERIOD is proximal to the SWI/SNF chromatin-remodeling complex (manuscript in preparation), we asked whether the complex plays a role in regulating these accessibility rhythms. We first conducted a systematic knockdown of individual SWI/SNF subunits specifically in clock neurons. While locomotor assays revealed circadian rhythm defects for multiple subunit knockdowns (Figure S1C, S1D), knockdown of Moira (MOR), a core subunit present in two copies per complex, produced the most severe phenotype: complete behavioral arrhythmicity under constant darkness (Figure 1B, 1C). This phenotype was recapitulated by expression of a dominant-negative version of BRM, the ATPase subunit of the complex^30^, suggesting that functional SWI/SNF activity is required for circadian rhythmicity (Figure S1B). Importantly, conditional adult-specific knockdown of MOR also resulted in strong behavioral defects, ruling out developmental artifacts (Figure S1B). To further validate these findings, we generated a tissue-specific *mor*-sgRNA knockout line (Figure S1A); these clock-specific knockout flies similarly exhibited arrhythmic locomotor activity, phenocopying the RNAi results (Figure 1B, 1C). Together, these data identify MOR as a critical SWI/SNF subunit essential for normal circadian behavior.

**Figure 1.**
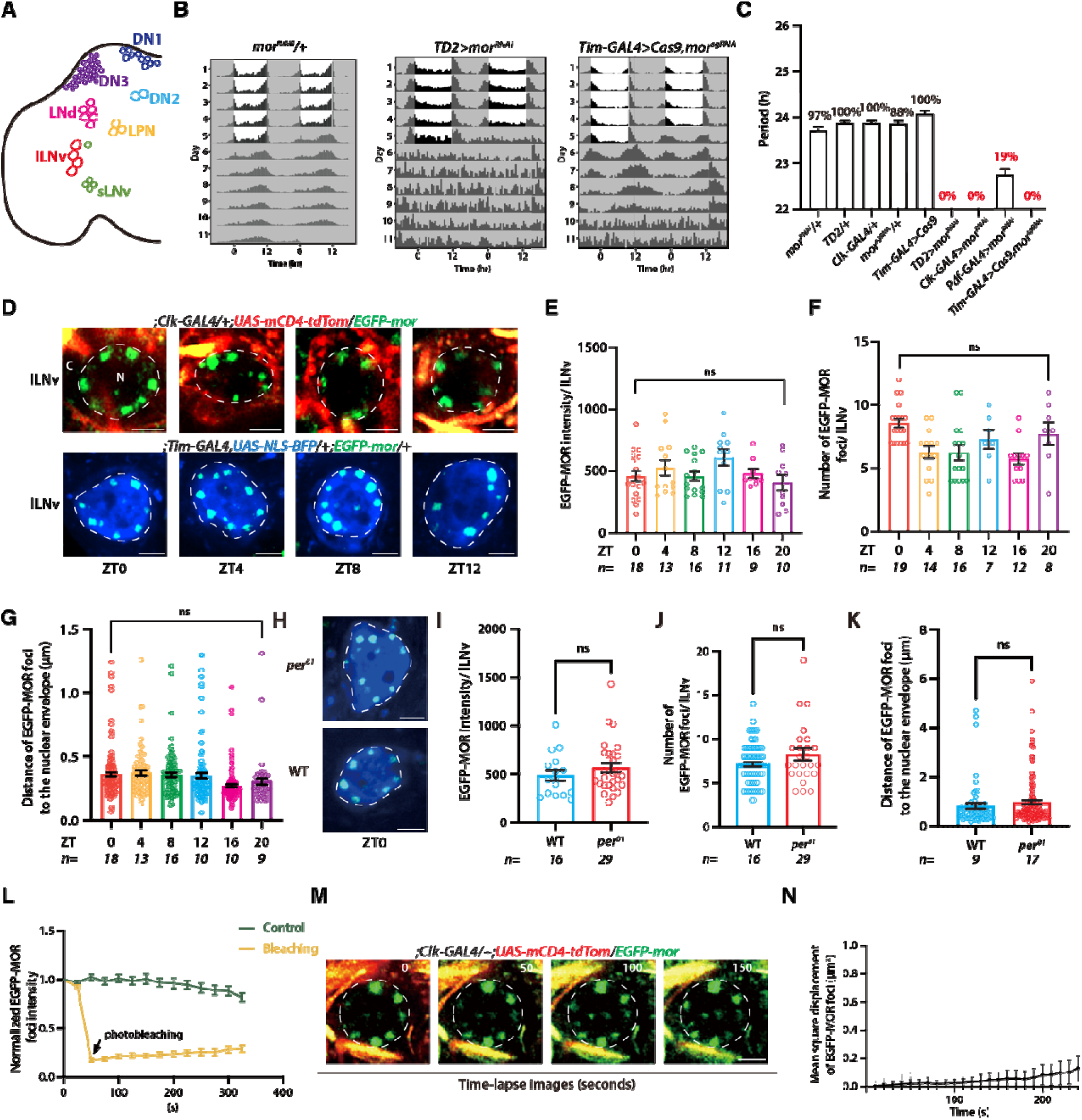
MOR is essential for circadian behavior and forms stable, foci anchored at the nuclear periphery in clock neurons. (A) Schematic of the clock neuron network in the adult *Drosophila* brain. (B) Representative actograms of control flies (*mor^RNAi^*/+) and flies with clock-neuron-specific knockdown (*TD2>mor^RNAi^*, TD2 corresponds to the *Tim-GAL4,UAS-Dicer2*) or knockout of mor (*Tim-GAL4>Cas9,mor^sgRNA^*) in LD and constant darkness (DD). (C) Quantification of rhythmicity for the genotypes shown in. MOR knockdown or knockout in clock neurons results in complete arrhythmicity (0% rhythmic). (D) Representative confocal images of endogenously tagged EGFP-MOR (green) in lLNv neurons (top, cell membrane marked by *Clk-GAL4>mCD4-tdTom*, red; bottom, nucleus marked by *Tim-GAL4>NLS-BFP*, blue) at different timepoints over the circadian cycle (ZT0, ZT6, ZT12, ZT18). MOR forms discrete nuclear foci (E-G) Quantification of EGFP-MOR foci properties in lLNv neurons at different timepoints. (E) number of foci per nucleus, and (F) Mean EGFP-MOR fluorescence intensity, (G) distance of foci to the nuclear envelope (µm). All parameters remain constant (ns, not significant) across the day, indicating that MOR foci abundance and localization are not oscillatory. (H-K) MOR foci formation is independent of the core clock. (H) Representative images of EGFP-MOR (green) in control and *per^01^* mutant neurons. (I-K) Quantification of (I) fluorescence intensity, (J) distance to the nuclear envelope, and (K) foci number shows no significant difference between control and *per^01^* mutants. (L-N) MOR foci exhibit stable and immobile biophysical properties. (L) Fluorescence Recovery After Photobleaching (FRAP) analysis of EGFP-MOR foci. Compared to an unbleached control (green), the photobleached focus (yellow) shows minimal recovery, indicating limited molecular exchange. (M) Time-lapse images of the FRAP experiment in (L). (N) Mean Square Displacement (MSD) analysis shows near-zero displacement, confirming that MOR foci are highly immobile within the nucleus. All scale bars here are 2µm.

To determine the mechanism by which MOR regulates the circadian clock, we first examined its spatial organization in *Drosophila* clock neurons. We generated a *mor* knock-in allele by fusing EGFP to the N-terminus using CRISPR/Cas9 (Figure S2A). We confirmed the functionality of the fusion protein in two ways: first, homozygous EGFP-MOR flies displayed control locomotor rhythms under both 12 h light-dark (LD 12:12) cycles and constant darkness (DD), with a ∼24 h free-running period; second, the EGFP-MOR allele rescued the behavioral arrhythmicity of *mor¹* mutant flies^26^ (Figure S2B). We then used this knock-in line to track MOR localization in individual clock neurons in live *Drosophila* brains. EGFP-MOR flies were entrained to LD 12:12 cycles, and brains were imaged every 4 h across the circadian cycle using Airyscan high-resolution confocal microscopy. We first focused on PDF-expressing ventral lateral neurons (LNvs), and found that EGFP-MOR forms distinct, puctate foci near the nuclear periphery (Figure 1D). This peripheral MOR foci organization was not unique to the LNvs but was observed across other clock neuron clusters (Figure S2F) and more broadly across neuronal cell types in the brain (Figure S2E). In the large LNvs (l-LNvs), we quantified ∼10 distinct MOR foci (Figure 1E, 1F) typically positioned within 0.5 μm of the nuclear envelope (Figure 1G). Unlike the highly dynamic PER foci^18^, MOR foci remained spatially stable with constant signal intensity throughout the circadian cycle. This organization persisted under temperature cycles (18°C/25°C), indicating that the subnuclear architecture of MOR is insensitive to environmental zeitgebers (Figure S2J, S2K).

To determine whether MOR foci formation depends on the core circadian clock, we analyzed *per*^01^ null mutants, which lack functional PERIOD protein and are molecularly and behaviorally arrhythmic^31^. MOR foci in *per*^01^ flies remained intact and were morphologically indistinguishable from those in control flies, indicating that MOR nuclear peripheral localization and foci formation are independent of PER and the molecular clock mechanism (Figure 1H-K).

Strikingly, we observed that MOR foci are confined exclusively to the nuclear periphery rather than being distributed throughout the nucleoplasm. If chromatin were uniformly distributed, one would expect MOR, which acts on chromatin, to be similarly dispersed throughout the nuclear volume. Instead, this restricted localization aligns closely with the unique architecture of the *Drosophila* nucleus: recent live-imaging studies show that chromatin is not uniformly distributed, but concentrated in a peripheral shell adjacent to the nuclear lamina, leaving the nuclear interior largely devoid of DNA^32^. Thus, MOR foci are positioned precisely at the subnuclear location where the genome resides in *Drosophila* cells.

To probe the biophysical nature of these foci, we performed fluorescence recovery after photobleaching (FRAP) on EGFP-MOR foci in neurons of intact fly brains. A defined region within individual foci was rapidly photobleached, and fluorescence recovery was monitored over time. MOR foci displayed minimal recovery over 5-10 minutes post-bleaching (Figure 1L), indicating highly restricted molecular exchange with the surrounding nucleoplasm. Such slow dynamics distinguish MOR foci from transient liquid condensates and are instead characteristic of stable nuclear bodies or “gel-like” scaffolds, similar to PML bodies or heterochromatin clusters, which show negligible recovery over tens of minutes^33, 34^. Consistent with this, 150 seconds of time-lapse imaging (Figure 1M, 1N) revealed that MOR foci are essentially immobile. Taken together, the persistent and immobile nature of MOR foci suggests they function as stable, anchored scaffolds at the nuclear periphery rather than dynamic liquid organelles.

Next, to investigate the mechanisms underlying MOR foci formation, we analyzed the MOR protein sequence using the intrinsic disorder prediction algorithm IUPred2A. This analysis revealed three distinct intrinsically disordered regions (IDRs) within the protein (Figure S3B), a structural feature frequently associated with proteins that undergo foci formation^35^. To test the self-assembling capacity of these regions, we purified recombinant full-length MOR and each IDR individually. Both the full-length protein and two of the IDR fragments readily formed micrometer-sized spherical foci *in vitro* under physiological salt conditions (Figure S3C-E), with IDR1 displaying the most robust assembly. These findings demonstrate that MOR possesses the intrinsic capacity for self-assembly mediated by its IDRs, suggesting a mechanism where IDR-mediated condensation initiates foci formation, which are then stabilized *in vivo* into stable architectural scaffolds at the nuclear periphery.Together, these findings establish Moira as an essential SWI/SNF subunit for circadian behavior and reveal that it is organized into discrete, stable foci at the nuclear periphery.

### Moira is required to generate rhythms in chromatin accessibility of clock-regulated genes

Given that MOR is essential for circadian behavior, we next asked whether it is required for circadian chromatin remodeling in clock neurons. To address this, we performed ATAC-seq on nuclei isolated from GFP-labeled clock neurons in control and experimental flies where MOR is knocked down specifically in clock neurons (*Clk>mor^RNAi^*) at two key timepoints: CT24 (repression phase) and CT36 (activation phase). Briefly, nuclei were sorted by FACS to enrich for GFP-positive clock cells, and ATAC-seq libraries were prepared using standard Tn5 tagmentation, followed by high-throughput sequencing and peak calling (Figure 2A).

**Figure 2.**
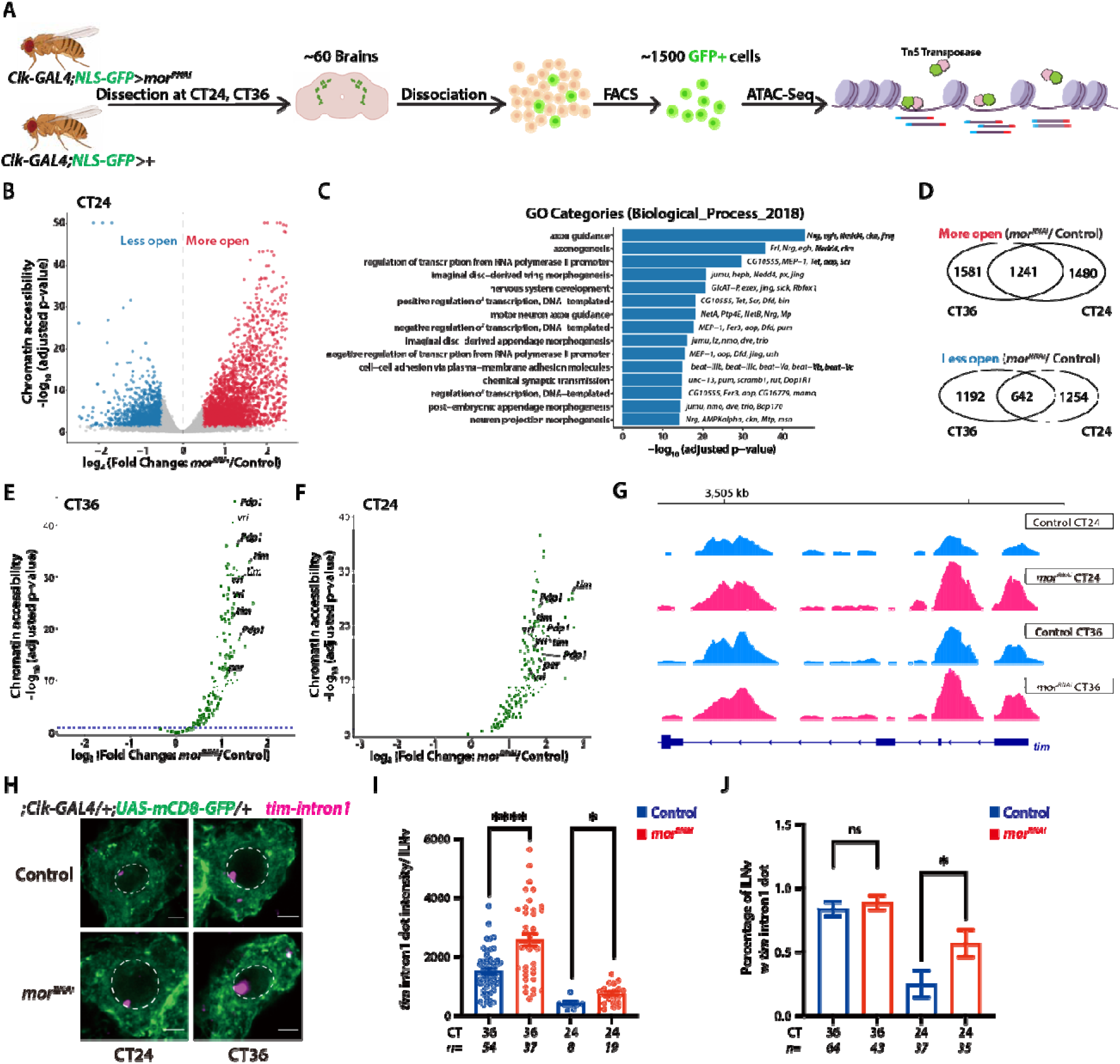
MOR is essential for rhythmic chromatin accessibility of clock-regulated genes. (A) Schematic diagram of the ATAC-seq experimental workflow. Nuclei from GFP-labeled clock neurons (*Clk-GAL4; UAS-NLS-GFP*) from either control or *mor^RNAi^* flies were isolated by Fluorescence-Activated Cell Sorting (FACS) at two circadian time points (CT24 and CT36). (B) Volcano plot showing differentially accessible regions in *mor^RNAi^* clock neurons compared to control at CT24. Genes with significantly increased accessibility (more open) are in red, and those with decreased accessibility (less open) are in blue. (C) Gene Ontology (GO) enrichment analysis for genes associated with differentially accessible regions identified at CT24. Enriched categories include regulation of transcription and various aspects of nervous system development. (D) Venn diagrams showing the overlap of differentially accessible regions (comparing *mor^RNAi^* vs. control) identified at CT24 and CT36. A large number of both more-open (top) and less-open (bottom) regions are shared between the two time points, indicating that the accessibility changes are largely constitutive and not time-of-day specific. (E-F) Volcano plots showing chromatin accessibility changes in *mor^RNAi^*neurons relative to control at CT36 (E) and CT24 (F). Green dots indicate genes that peak at CT36 in control (e.g., Clk). Notably, CT36-peaking genes display constitutively elevated accessibility in *mor^RNAi^* flies (log₂FoldChange > 0) both in CT36 and CT24. (G) Representative ATAC signal pile-up tracks at *tim* loci. Signal is normalized to total number of reads for each sample and therefore is comparable among samples. Four different replicates are overlaid. *mor^RNAi^* flies show stronger ATAC signal in *timeless* (*tim)* loci compared to the control group both in CT36 and CT24. (H) Representative images of HCR-FISH for *tim* nascent transcripts (*tim* intron 1, the first intron of *tim* gene, magenta) in lLNv neurons (cytoplasm marked by *Clk-GAL4>mCD8-GFP*, green) from control and *mor^RNAi^* flies at CT24 and CT36. (I) Quantification of *tim* intron signal intensity in lLNvs. Nascent transcription of *tim* is significantly elevated in *mor^RNAi^* flies (red) compared to control (blue) at both circadian time points (CT24 and CT36). (J) Quantification of the percentage of lLNvs positive for *tim* intron signal. The proportion of cells actively transcribing *tim* is significantly higher in *mor^RNAi^* flies at both time points. Statistics: Error bars represent mean ± SEM. *p < 0.05, ***p < 0.0001, determined by t-test.

We observed widespread alterations in chromatin accessibility in mor knockdown flies compared to controls, with thousands of loci showing significant differences (Figure 2B, S4A, 2D). Consistent with the established role of SWI/SNF as a chromatin remodeler, we found that MOR modulates chromatin accessibility bidirectionally across the genome, with substantial overlap between upregulated and downregulated regions at both time points. Gene ontology analysis of differentially accessible regions revealed significant enrichment for genes involved in transcriptional regulation, nervous system development, and appendage morphogenesis (Figure 2C, S4B), highlighting the broad regulatory impact of MOR-dependent chromatin remodeling in *Drosophila* clock neurons.

We next focused on 116 “clock-regulated” loci—including core genes like *per* and *tim*—that exhibit rhythmic accessibility in control flies, cycling between a closed state at CT24 and an open state at CT36^6^. In striking contrast to its bidirectional role globally, MOR appears to play a restrictive role at these clock loci. Knockdown of *mor* completely abolished accessibility rhythms; instead of cycling, these clock-regulated loci remained constitutively hyper-accessible at both timepoints (Figure 2E, 2F). To gauge the magnitude of this deregulation, we compared *mor* knockdown flies to *per*[*¹* null mutants, which also exhibit elevated accessibility due to the loss of the core repressor. Remarkably, we found that chromatin accessibility in *mor* knockdown neurons was significantly higher than in *per*[*¹* mutants (Figure S4C, S4D). Representative ATAC-seq tracks at the *tim* locus illustrate this hierarchy: in the absence of MOR, promoter and enhancer regions exhibit hyper-accessible peaks that consistently exceed levels observed in both control and *per*[*¹* backgrounds (Figure 2G, S4E).

To determine the transcriptional consequences of this constitutively open chromatin, we examined nascent transcription of the core clock gene *timeless* (*tim*) using hybridization chain reaction fluorescence in situ hybridization (HCR-FISH). Using intron-specific probes to detect nascent RNA in intact tissues^6^, we observed significantly elevated *tim* transcriptional activity in *mor* knockdown neurons at both the repression (CT24) and activation (CT36) phases (Figure 2H-J). This transcriptional hyperactivity mirrors the hyper-accessible chromatin state observed in our ATAC-seq data, confirming that the loss of accessibility rhythms leads to deregulated transcriptional output. Collectively, these findings demonstrate that MOR is essential for maintaining chromatin accessibility rhythms over the circadian cycle; without it, clock loci default to a constitutively open and transcriptionally active state. Thus, MOR acts as a key architect of the repressive chromatin environment required for molecular clock function.

### Moira-dependent chromatin remodeling is required to translate PER/TIM protein rhythms into rhythmic gene expression

We next determined how these MOR-dependent chromatin accessibility changes translate to gene expression. To address this, we performed RNA-seq on FACS-sorted clock neurons from control and *mor^RNAi^*flies collected in constant darkness at six-hour intervals (CT24-CT42). We compared gene expression between control and knockdown flies (Figure 3A, S5B-D) and observed widespread transcriptional dysregulation. Both oscillating and non-oscillating genes displayed significant dysregulation, suggesting that MOR regulates not only circadian targets but also broader transcriptional programs, consistent with the global changes in chromatin accessibility observed in Figure 2. Gene Ontology (GO) analysis of differentially expressed genes revealed enrichment for chromatin organization, transcriptional control, and neural signaling, in line with the pleiotropic functions of SWI/SNF complexes (Figure 3B, S5E-G).

**Figure 3.**
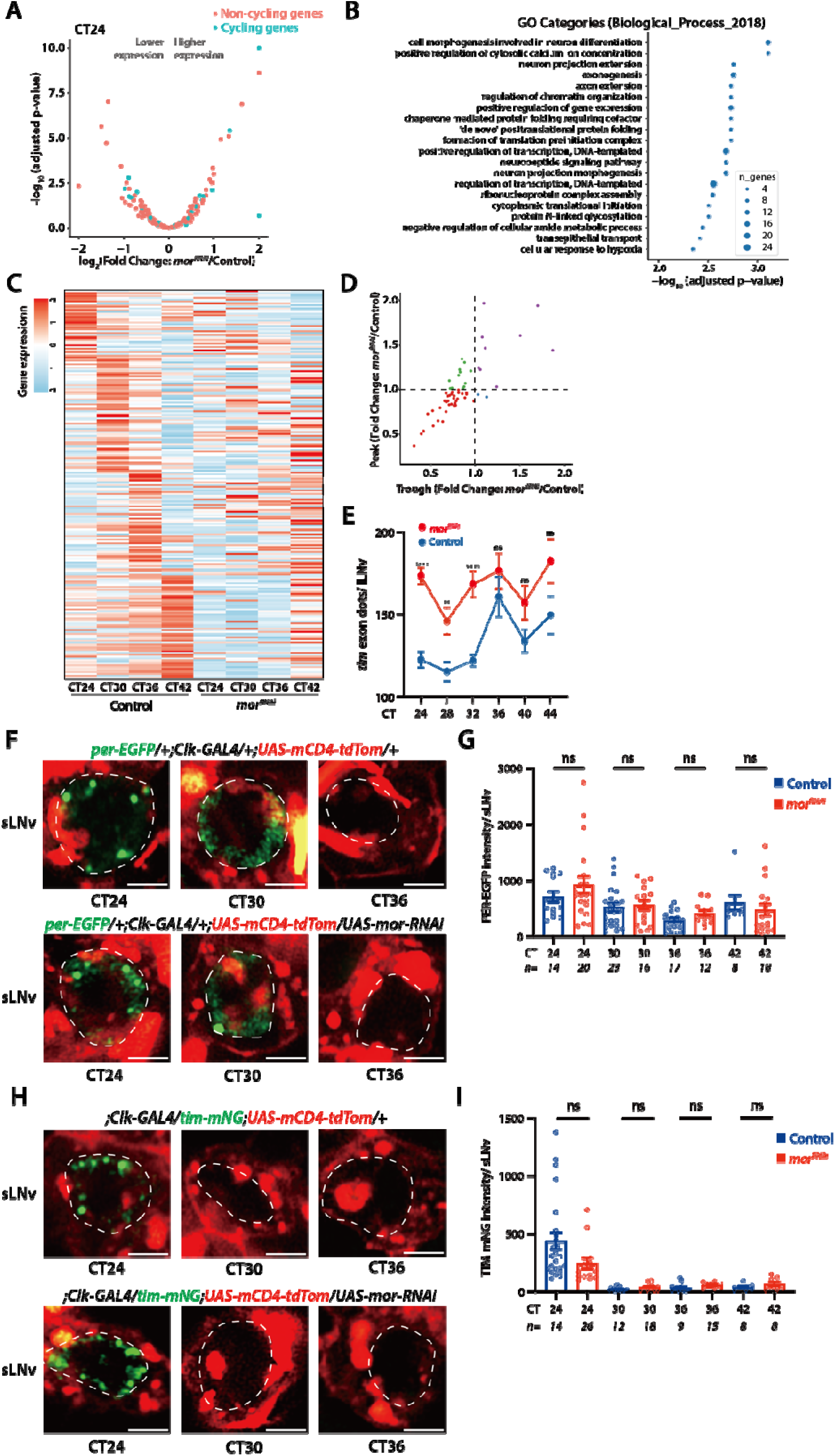
MOR knockdown disrupts rhythmic transcription of clock-controlled genes but does not affect the oscillation of core clock proteins PER and TIM. (A) Volcano plot of differential gene expression in *mor^RNAi^* vs. control at CT24. Both non-cycling (red) and cycling (blue) genes show significant expression changes, indicating broad transcriptional dysregulation. (B) Gene Ontology (GO) analysis of genes differentially expressed in *mor^RNAi^* flies at CT24, showing enrichment for categories related to chromatin organization and transcriptional regulation. (C) Heatmap of oscillating transcripts identified by RNA-seq in FACS-sorted clock neurons from control and *mor^RNAi^* flies at four circadian time points CT24, CT30, CT36, CT42. Rhythmic gene expression is largely abolished in neurons. (D) Scatter plot comparing the peak fold change and trough fold change of cycling genes in *mor^RNAi^* vs. control. Most genes (red dots) show reduced amplitude in the knockdown group. (E) Quantification of *tim* mature mRNA (exon dots) by HCR-FISH in lLNv neurons. In *mor^RNAi^* flies (red), the rhythmic expression of both genes is significantly altered and dampened compared to control (blue). (F) Representative live-imaging of endogenous PER-EGFP (green) in sLNv neurons (cytoplasm marked by Clk-GAL4>mCD4-tdTom, red) in control (top) and *mor^RNAi^* (bottom) flies at CT24, CT30, and CT36. (G) Quantification of PER-EGFP intensity from (F). There is no significant difference in PER protein levels or oscillation between Control (blue) and mor^RNAi^ (red) flies. (H) Representative live-imaging of endogenous TIM-mNeonGreen (TIM-mNG, green) in sLNv neurons in Control (top) and *mor^RNAi^* (bottom) flies. (I) Quantification of TIM-mNG intensity from (H). Similar to PER, there is no significant difference in TIM protein levels or oscillation between control and *mor^RNAi^*flies. Statistics: Error bars represent mean ± SEM. *p < 0.05, ***p < 0.0001, determined by t-test.

In control flies, over 200 genes exhibited rhythmic expression, including well-characterized clock-controlled genes such as *per*, *tim*, *vrille*, and *pdp1*. In contrast, mor knockdown caused a widespread disruption of rhythmic gene expression: only ∼50 genes retained detectable cycling, and even these displayed markedly altered phase and dampened amplitude (Figure 3C-D, S5A). These findings indicate that MOR is broadly required to maintain the transcriptional rhythmicity of the circadian program. To validate these results at single-cell resolution, we performed HCR-FISH using exon-targeting probes for *tim*. Consistent with the RNA-seq data, *tim* mRNA levels were elevated and displayed dampened rhythmicity in *mor^RNAi^* flies compared to controls (Figure 3E), directly mirroring the hyper-accessible chromatin state observed at the *tim* locus.

We next asked whether these transcriptional defects propagated to the protein level. Using live imaging of flies endogenously expressing PER-EGFP^9^ or TIM-mNeonGreen^10^, we assessed protein dynamics in the small ventrolateral neurons (sLNvs). Surprisingly, despite the disruption of mRNA rhythms, we observed no detectable difference in PER or TIM protein levels, subnuclear localization, or oscillation phase between control and *mor^RNAi^* flies (Figure 3F-I). PER and TIM nuclear foci remained intact and oscillated in intensity indistinguishably from controls (Figure S6).This striking dissociation between disrupted mRNA and robust protein oscillations is likely maintained by post-transcriptional mechanisms such as translational control (e.g., via ATX2^36, 37^) or post-translational stability^38^. Crucially, the persistence of PER/TIM protein rhythms in behaviorally arrhythmic flies demonstrates that MOR loss functionally uncouples the molecular oscillator from its genomic output. We conclude that MOR acts as a required gatekeeper of the circadian feedback loop: by establishing a permissive yet restrained chromatin landscape at clock loci, MOR ensures that rhythmic protein activity is faithfully translated into rhythmic transcriptional repression.

### Clock-regulated genes are organized into peripheral hubs at the nuclear periphery

The static and peripheral nature of MOR foci, combined with their role in repressing clock-regulated chromatin, led us to hypothesize that clock-regulated genes are localized to these foci, forming stable architectural compartments at the nuclear periphery. To test this, we used DNA fluorescence in situ hybridization (DNA-FISH) to examine the subnuclear localization of the core clock gene *period* (*per*). We focused on DN1 dorsal clock neurons for these experiments, as their superficial position provides superior optical accessibility for high-resolution nuclear imaging. Using an NLS-GFP line to define the nuclear boundary, we found that the *per* locus was constitutively positioned at the nuclear periphery at both ZT0 and ZT12 (Figure 4A), placing it in close proximity to the nuclear envelope where MOR foci reside. Consistent with the established pairing of homologous chromosomes in *Drosophila* somatic cells^39^, we detected a single per FISH signal per nucleus.

**Figure 4.**
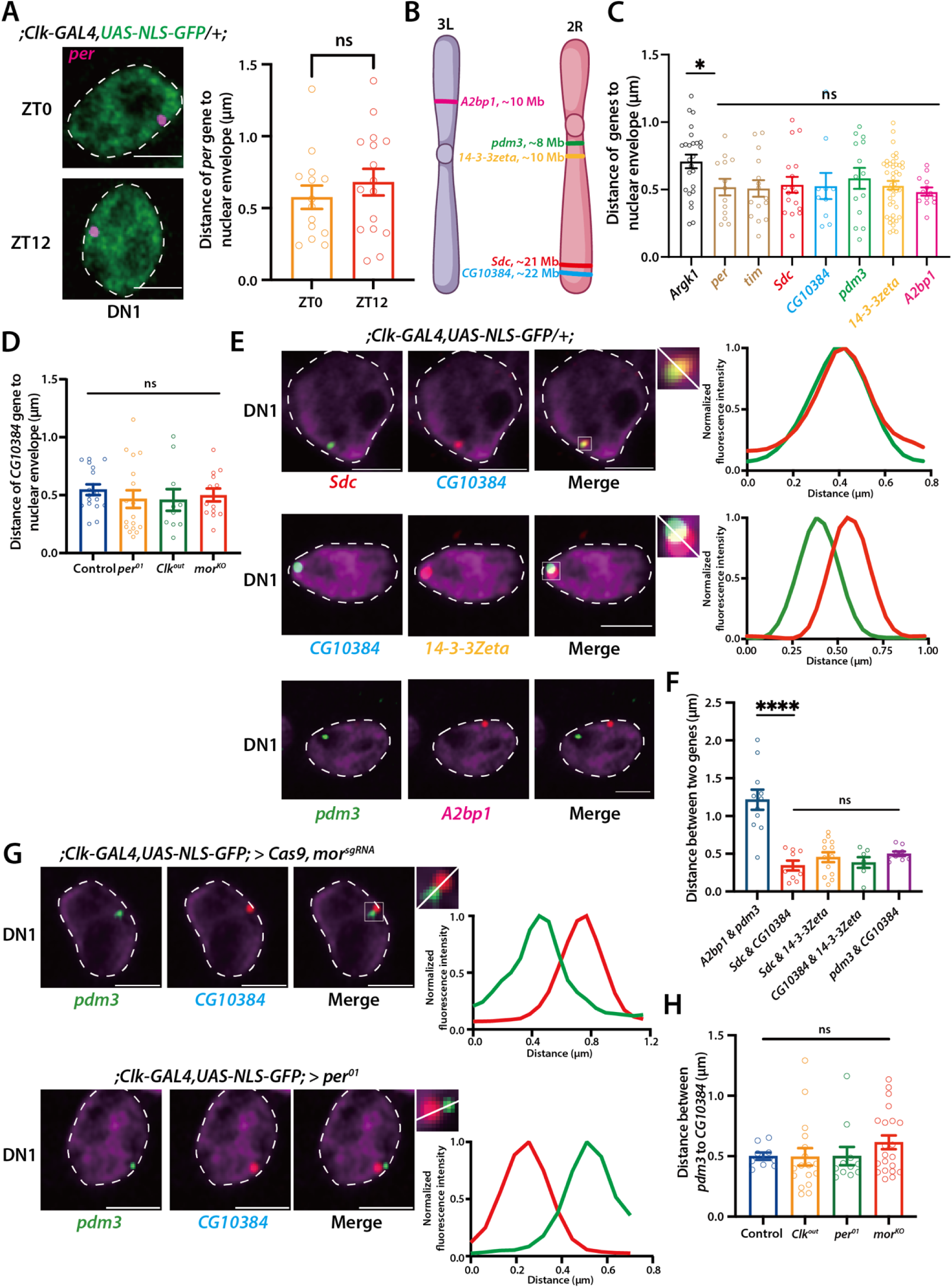
Clock-regulated genes are organized into pre-existing, static hubs at the nuclear periphery, independent of the circadian clock and MOR. (A) Representative DNA-FISH images of the *per* locus (magenta) in clock neurons (nucleus labeled by NLS-GFP, green) at ZT0 and ZT12. The graph (right) quantifies the distance from the *per* locus to the nuclear envelope, showing it is constitutively anchored at the periphery with no significant difference between time points. (B) Schematic of *Drosophila* chromosomes 2R and 3L, showing the genomic locations of the five clock-regulated genes (*pdm3*, *14-3-3*ζ, *Sdc*, *CG10384*, *A2bp1*) targeted for DNA-FISH. (C) Quantification of the distance from various gene loci to the nuclear envelope. The housekeeping gene *Argk1* shows a random, nucleoplasmic distribution (larger distance), while all tested clock-regulated genes (*per*, *tim*, *pdm3*, *14-3-3*ζ, *Sdc*, *CG10384*, *A2bp1*) are significantly anchored at the nuclear periphery (*p < 0.05). (D) Quantification of the *CG10384* locus distance to the nuclear envelope in control, core clock mutants (*per^01^*, *Clk^out^*), and *mor* knockout (*Clk-GAL4>Cas9,mor^sgRNA^*) flies. Peripheral anchoring is maintained. (E) Representative DNA-FISH images show intra-chromosomal clustering. Gene loci on the same chromosome are spatially clustered (*Sdc & CG10384*, *CG10384 & 14-3-3*ζ), as shown in merge insets and corresponding line-intensity plots (right), regardless of their relative positions along the chromosome. Loci on different chromosomes are spatially segregated (*pdm3 & A2bp1*). (F) Quantification of the 3D distance between pairs of gene loci. Inter-chromosomal pairs (*A2bp1* & *pdm3*) are significantly more segregated (larger distance) than intra-chromosomal pairs, which are tightly clustered (****p < 0.0001). (G) Representative DNA-FISH images show that in both *mor* knockout and *per^01^* backgrounds, the *pdm3* and *CG10384* loci co-localize at the nuclear periphery, suggesting that this subcellular organization is not dependent on clock function or MOR. (H) Quantification of the distance between *pdm3* & *CG10384* locus in control, core clock mutants (*per^01^*, *Clk^out^*), and *mor* knockout neurons. Clustering is maintained in all conditions. Statistics: Error bars represent mean ± SEM. *p < 0.05, ***p < 0.0001, determined by t-test.

To determine whether this organization extends beyond per, we designed DNA-FISH probes targeting five additional clock-regulated genes identified from a published RNA-seq dataset^40^: *pdm3*, *A2bp1*, *Sdc*, *CG10384*, and *14-3-3*ζ. These genes are rhythmically expressed specifically in DN1 neurons, with peak expression during the activation phase, and are distributed across two major chromosomes (3L and 2R) (Figure 4B). Strikingly, DNA-FISH revealed that, similar to *per*, all five clock-regulated genes were positioned close to the nuclear periphery, whereas the housekeeping gene *Argk1* exhibited a dispersed, nucleoplasmic distribution (Figure 4C). Thus, peripheral positioning appears to be a shared and non-random feature of clock-regulated loci.

We next asked how dynamic this spatial architecture is and whether it depends on the circadian clock or MOR. Intriguingly, the peripheral localization of these genes was maintained regardless of time of day (LD vs. DD) and persisted in *per*^O^*^¹^*, *Clk^jrk^* mutants, and *mor^RNAi^* conditions (Figure 4D, Figure S7A). These results indicate that perinuclear positioning of clock-regulated genes is established independently of both the core molecular clock and MOR.

This shared perinuclear anchoring prompted us to examine how these genes are arranged relative to one another in three-dimensional space. We found that clock-regulated genes located on different chromosomes (e.g., *A2bp1* on 3L and *pdm3* on 2R) were spatially segregated, consistent with the fact that different chromosomes occupy distinct chromosome territories^41^. By contrast, clock-regulated genes located on the same chromosome were frequently clustered together, even when separated by large linear genomic distances (Figure 4E-F). To quantify this organization, we examined specific loci on chromosome 2R spanning a range of linear distances. As expected, *Sdc* and *CG10384*, which are separated by only ∼1 Mb (at 21 Mb and 22 Mb, respectively), exhibited high spatial colocalization. Remarkable, however, we also observed frequent clustering between loci separated by over 10 Mb, such as *CG10384* and *14-3-3*ζ. These observations are consistent with higher-order chromatin folding that brings distant clock-regulated loci into close physical proximity, organizing them into discrete intra-chromosomal “hubs” within the nucleus.

Crucially, this clustered perinuclear architecture did not require either MOR or the core molecular clock: these gene hubs remained intact in both *mor* knockdown and *per*[*¹* neurons (Figure 4G-H). Moreover, this organization appears to be a constitutive feature of clock neuron nuclear architecture rather than a consequence of active transcription. Although these five genes are expressed in DN1a neurons, they remained similarly clustered at the nuclear periphery in lLNv neurons, where they are transcriptionally silent (Figure S7C-D).

Taken together, these findings reveal that clock-regulated genes are organized into stable “hubs” at the nuclear periphery. Importantly, this spatial architecture is established independently of MOR or the circadian clock. Given that MOR forms foci in this same peripheral compartment, this raises a critical mechanistic question: do these genes physically colocalize with the MOR-containing SWI/SNF foci to enable rhythmic regulation?

### MOR forms a stable nuclear scaffold that gates dynamic circadian repression

To test this idea, we sought to visualize both the gene (*tim*) and the MOR protein simultaneously. Initially, we attempted to combine DNA-FISH with protein imaging. However, despite extensive optimization, MOR fluorescence was completely lost during the high-temperature denaturation steps required for DNA-FISH, likely due to disruption of the fluorescent protein structure. To overcome this limitation, we implemented intron RNA-FISH using hybridization chain reaction (HCR-FISH). Because most introns are co-transcriptionally spliced, probes targeting intronic regions hybridize to nascent RNA still tethered to the chromatin. This approach provides a precise proxy for the genomic position of the gene itself^6, 42^, while using mild conditions that preserve protein fluorescence.

We combined EGFP-MOR labeling with intron-specific HCR-FISH probes against *timeless* (*tim*) and the voltage-gated potassium channel *shaker* (*Sh)*, which served as a non-clock-regulated negative control. Confocal imaging at ZT12 (peak transcription) revealed that *tim* transcription sites frequently colocalized with MOR foci, whereas *Sh* signals showed minimal spatial proximity (Figure 5A). Quantification confirmed that a high percentage of *tim* transcriptional loci were embedded within MOR foci, while *Sh* loci were randomly distributed (Figure 5B). Because intron-specific probes detect nascent RNA at the gene transcription site, these experiments confirm that the *tim* locus is physically localized within the MOR compartment.

**Figure 5.**
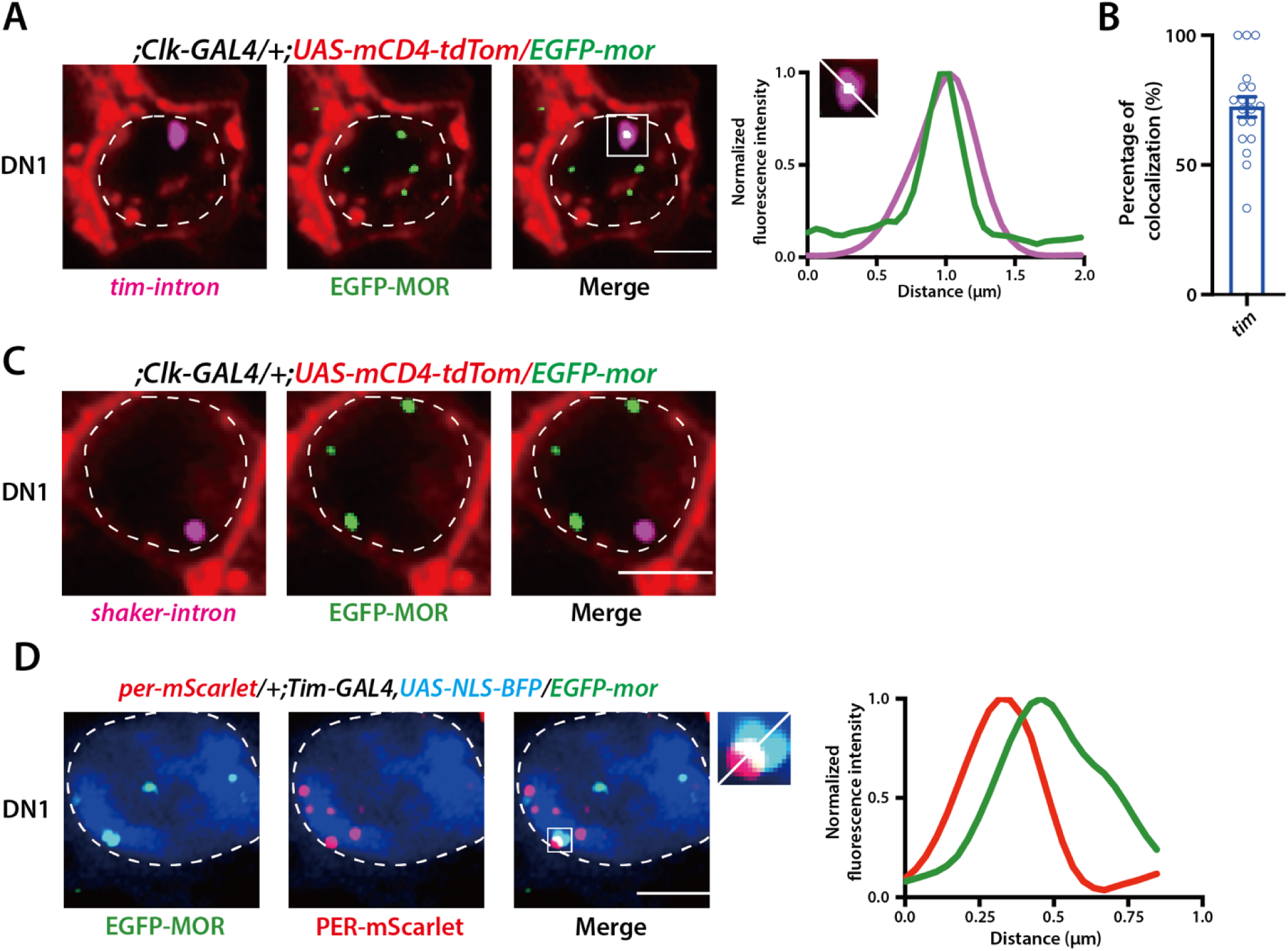
MOR co-localizes with clock-regulated gene hubs and is adjacent to PER. (A) Representative images showing co-localization of EGFP-MOR (green foci) with active transcription sites (magenta) of *tim*, detected by intron-specific HCR-FISH. The negative control gene shaker (bottom) shows no co-localization. Corresponding line-intensity plots (right) confirm the spatial overlap for *tim*. (B) Quantification of the percentage of colocalization between MOR foci and the indicated gene loci from (A). *tim* transcription sites show significantly higher association with MOR compared to shaker (*p < 0.05). (C) Dual-color live imaging of EGFP-MOR (green) and the core clock protein PER-mScarlet (red) in DN1 (nucleus labeled with *Tim>NLS-BFP*, blue). Merge insets and line-intensity plots (right) show that MOR and PER signals are in close proximity.

Finally, we investigated how the dynamic core repressor PER interfaces with this stable MOR scaffold. Our previous work demonstrated that during the repression phase, PER accumulates in discrete nuclear foci away from the clock loci^9, 10^. To explore the spatial relationship between these dynamic PER foci and the static MOR foci, we performed dual-color imaging of EGFP-MOR and PER-mScarlet in clock neurons during the repression phase (ZT0). We observed that MOR and PER signals frequently appear in close spatial proximity but do not overlap, suggesting that these proteins occupy adjacent but distinct subnuclear compartments (Figure 5C). This reveals that clock genes are anchored within the MOR chromatin scaffold, physically distinct from the adjacent foci where PER accumulates.

### Moira requires Otefin for perinuclear foci formation

We next asked whether specific protein interactions are required for MOR foci to be anchored at the nuclear envelope *in vivo*. We hypothesized that inner nuclear membrane proteins might contribute to their assembly or localization. Interestingly, previous proteomics studies have shown that the SWI/SNF complex interacts with inner nuclear envelope proteins, specifically LEM-domain proteins such as Emerin, the mammalian homolog of *Drosophila* Otefin^29, 43^. LEM-domain proteins (e.g., LAP2, Emerin, MAN1) are inner nuclear membrane components that physically connect the nuclear lamina to chromatin^29^. These attributes make LEM-domain proteins attractive candidates for organizing MOR foci; specifically, we asked—do they provide the tether that anchors MOR to perinuclear chromatin?

To test this possibility, we examined *Otefin* null mutant^44^ flies and found that they exhibited severe circadian arrhythmia (Figure 6A). Otefin is a LEM-domain protein known to cooperate with the Barrier-to-Autointegration Factor (BAF) to anchor chromatin to the inner nuclear membrane and maintain repressed heterochromatin at the nuclear periphery^29, 43^. To directly examine the subcellular localization of Otefin, we generated Ote-V5–EGFP flies using CRISPR/Cas9-mediated tagging at the endogenous *otefin* locus (Figure S8A). Immunostaining with an anti-V5 antibody combined with DAPI nuclear staining revealed that Otefin forms discrete puncta along the nuclear envelope, typically displaying ∼10 foci per nucleus (Figure 6E). This punctate pattern contrasts with the canonical view of LEM-domain proteins, which are typically described as being uniformly distributed along the inner nuclear membrane. Notably, these discrete Otefin foci were frequently positioned adjacent to the *tim* gene locus, as visualized by combined protein imaging and HCR-FISH (Figure 6F). We note that simultaneous visualization of Otefin and MOR proteins was precluded in these experiments, as both tagged alleles share identical V5 and EGFP epitopes.

**Figure 6.**
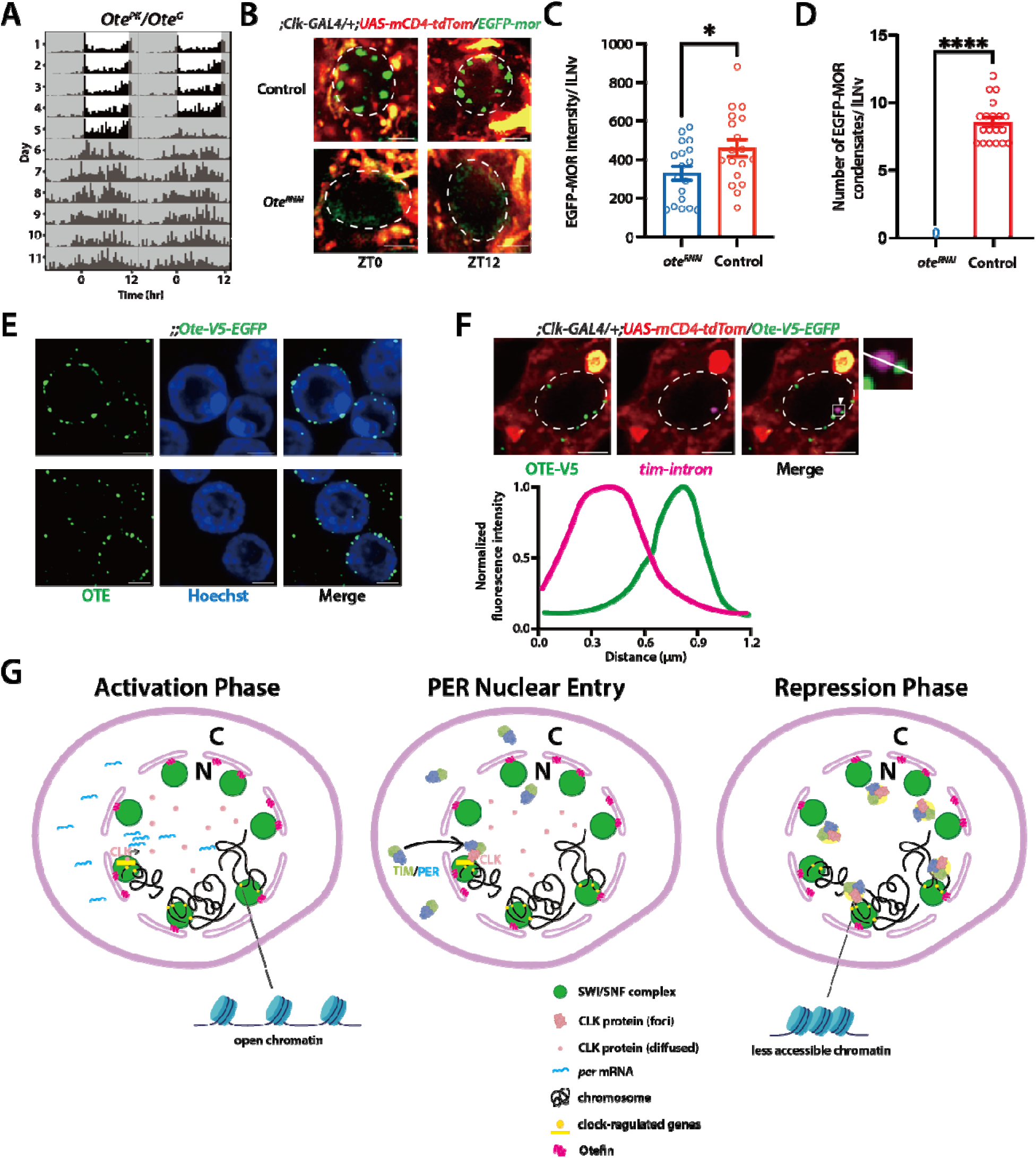
MOR in vivo anchoring at the nuclear periphery is dependent on the LEM-domain protein Otefin, acting as a stable scaffold to gate rhythmic transcription. (A) Representative actogram of flies with Otefin null mutant in LD and constant darkness (DD), which exhibit a distinct rhythmic phenotype. (B) Representative confocal images of EGFP-MOR (green) localization in lLNv neurons (membrane marked by *Clk-GAL4>mCD4-tdTom*, red) from Control and Ote-RNAi flies at ZT0 and ZT12. MOR forms distinct foci in control neurons but becomes diffuse throughout the nuclear periphery upon Otefin knockdown. (C, D) Quantification of EGFP-MOR foci in lLNv neurons at ZT0. Compared to controls, *Ote^RNAi^* neurons show a mild reduction in (C) EGFP-MOR intensity per nucleus and significant reduction in (D) the number of discrete foci. (E) Confocal images showing the localization of endogenous Otefin (OTE, green) in *Drosophila* neurons stained by V5 antibody staining. OTE forms discrete puncta along the nuclear envelope, visualized by Hoechst staining (blue). (F) Representative images showing the spatial relationship between OTE-V5 (green) and the *tim* intron1 locus via HCR-FISH in clock neurons. White arrowheads indicate instances of close proximity. (G) Proposed model for MOR’s role as an architectural gating platform. Clock-regulated genes are positioned close to the nuclear periphery and clustered into hubs. The MOR (SWI/SNF) complex co-localizes with these hubs. During the activation phase, CLK proteins are present in a diffuse nuclear distribution and bind to clock-regulated genes, promoting transcription in a more accessible chromatin environment. Later, newly synthesized PER protein subsequently forms a heterodimer with TIM, translocates into the nucleus, and associates with CLK. In the repression phase, the PER/TIM/CLK complex forms condensate-like assemblies that position away from the gene bodies (and MOR foci). Statistics: Error bars represent mean ± SEM. *p < 0.05, ***p < 0.0001, determined by t-test.

The perinuclear distribution of Otefin strikingly resembled that of MOR foci, prompting us to investigate whether MOR foci formation relies on Otefin. In *Otefin*-depleted clock neurons, MOR failed to form discrete foci and instead appeared dispersed throughout the nucleoplasm (Figure 6B, 6D), even though global protein levels remained comparable to controls (Figure 6C). Thus, Otefin is required for the assembly and anchoring of MOR foci at the nuclear envelope. Together, these findings identify Otefin as the critical architectural anchor that secures the MOR-containing SWI/SNF complex to the nuclear periphery, enabling the spatial organization of the circadian genome.

## DISCUSSION

Based on these findings, we propose a model wherein MOR functions as a stable architectural component essential for gating rhythmic transcription (Figure 6G). Our data establishes that clock-regulated genes are not randomly distributed but are organized into clustered “hubs” anchored at the nuclear periphery. MOR-containing SWI/SNF complexes co-localize with these gene hubs, tethered to the nuclear envelope by the LEM-domain protein Otefin. Crucially, MOR is required to translate this static architecture into dynamic chromatin states. Specifically, we found that MOR is essential for generating rhythms in chromatin accessibility; without MOR, these loci fail to cycle between open and closed states. During the repression phase, the dynamic PER/TIM complex enters the nucleus and transiently associates with these clock loci and sequester the CLK/CYC complex away from chromatin. We propose that the MOR scaffold provides the necessary physical substrate for PER/TIM to recruit additional repressors, enforcing the maximal chromatin compaction required to silence CLK/CYC-driven transcription. Conversely, in the absence of MOR, this functional platform is lost. Although the gene hubs remain at the periphery, the chromatin fails to oscillate; consequently, even as PER/TIM complexes accumulate, they fail to implement robust silencing. Thus, robust circadian rhythmicity emerges from the integration of dynamic biochemical signaling with nuclear architecture. MOR establishes a stable physical platform that licenses the chromatin for rhythmic accessibility, enabling the core clock machinery to drive high-amplitude cycles of gene expression.

### Nuclear topology defines the circadian landscape

Our live imaging of endogenously tagged MOR reveals, for the first time, that it organizes into stable, discrete foci at the nuclear periphery of *Drosophila* clock neurons. While previous biochemical characterizations established that the intrinsically disordered regions (IDRs) of SWI/SNF subunits can drive phase separation *in vitro*^45^, our high-resolution imaging of endogenously tagged EGFP-MOR provides the first direct evidence of these higher-order assemblies within their native physiological context. Notably, unlike the dynamic liquid behaviors often observed in test-tube reconstitutions^45^, we demonstrate that native MOR foci function as stable scaffolds at the nuclear periphery in *Drosophila* cells. By visualizing these structures in intact *Drosophila* brains, we establish that MOR foci are not merely potential biochemical states but are fundamental architectural features of the nuclear periphery, positioned precisely to organize the circadian genome. Furthermore, we observed that MOR organizes into similar peripheral foci across various other neuronal groups in the brain. This suggests an intriguing possibility: that genes regulated by the SWI/SNF complex are systematically organized into these peripheral hubs.

A fundamental insight from this work is that the circadian genome operates within a fixed nuclear topology: clock genes are clustered into peripheral hubs. This spatial organization is intrinsic to the identity of the clock neuron: it persists independently of the circadian cycle, transcriptional activity, or the MOR scaffold. Within this static spatial framework, MOR performs a critical chromatin-remodeling function. In mor-depleted neurons, chromatin accessibility at clock loci is uniformly high, and *tim* nascent transcription remains elevated throughout the day. This indicates that MOR acts as a tonic repressor: it lowers basal chromatin accessibility at clock loci, providing the base state upon which the core clock machinery imposes rhythmic opening and closing. Without this MOR-mediated constraint, the clock operates on a constitutively permissive template, blunting the amplitude of the entire cycle.

This finding resonates with emerging evidence challenging the canonical view of SWI/SNF as solely a transcriptional activator. For instance, in mammalian embryonic stem cells, SWI/SNF complexes act as tonic repressors that condition the genome for LIF/STAT3 signaling^46^. Our results extend this paradigm to circadian regulation: MOR-containing SWI/SNF complexes at peripheral hubs establish the baseline nucleosome organization upon which PER/TIM-mediated repression and CLK/CYC-mediated activation are superimposed.

### Decoupling clock protein rhythms from transcriptional output

The necessity of this scaffold is most clearly demonstrated by the decoupling of clock protein rhythms from transcriptional output in mor knockdown flies. We observed that PER and TIM proteins continue to oscillate, enter the nucleus, and form nuclear foci with near-normal dynamics, yet they fail to drive rhythmic gene expression, suggesting that the core oscillator requires a specific chromatin interface to function. The stable MOR scaffold functions as an architectural gate that determines where and how the transient PER signal is interpreted. When the scaffold is intact, PER docking triggers chromatin compaction and silencing; when the scaffold is absent, the PER signal is effectively ignored by the genome.

In summary, we demonstrate that *Drosophila* circadian transcription is executed within a stereotypical peripheral nuclear architecture, where clock-regulated genes are organized into spatial hubs. Anchored by Otefin, MOR-containing SWI/SNF complexes localize to these sites to establish a tonic repressive chromatin state. MOR gates the activity of the core clock machinery, ensuring that the PER/TIM protein oscillations are translated into robust transcriptional silencing. This reliance on a stable physical platform to interpret dynamic temporal signals likely represents a general principle for organizing spatiotemporal gene expression.

## MATERIALS AND METHODS

### Fly stocks

Flies were raised on standard cornmeal/yeast media and maintained at 25°C under a 12h:12h light-dark (LD) schedule. For all experiments, flies were entrained to LD cycles for 5-7 days. The initiation of the light phase is defined as Zeitgeber Time 0 (ZT0), and the initiation of the dark period is ZT12. For constant darkness (DD) experiments, flies were entrained and then released into complete darkness. Circadian Time (CT) 0 marks the time when lights would have turned on.

The following fly lines were used: *Clk-GAL4*, *Tim-GAL4*, *Tim-GAL4*; *UAS-Dicer2*, *per*^01^, *Clk^Jrk^*, *UAS-CD4-tdTomato*, *UAS-NLS-BFP*^10^, *Clk-GAL4; tubgal80ts* and endogenously tagged PER-EGFP^9^, Tim-mNeonGreen^10^, PER-mScarlet^10^. RNAi lines (stocks numbers are provided in the data tables) were obtained from the Bloomington Drosophila Stock Center (BDSC) or the Vienna Drosophila Resource Center (VDRC).

### Generation of CRISPR-Cas9 knock-in lines

The endogenous EGFP-MOR knock-in line was generated using CRISPR/Cas9-mediated homologous recombination, following protocols similar to those described in a published report^9^. Briefly, two guide RNAs (gRNAs) were designed to target the N-terminal region of the mor locus at the start codon (ATG). A donor plasmid was constructed containing the EGFP sequence flanked by ∼1kb homology arms corresponding to the regions upstream and downstream of the target site. The donor plasmid and gRNA plasmids were injected into vasa-Cas9 embryos.

Progeny (F1) were screened by single-fly PCR to identify positive insertions. Positive flies were backcrossed to a w118 background, and homozygous stocks were established. The final EGFP-MOR knock-in allele was confirmed by PCR and Sanger sequencing. Functionality was validated through behavioral rescue of the *mor^[1]^* null mutant by confirming normal per mRNA oscillations and locomotion behavior.

### Generation of CRISPR-Cas9 knockout lines

Tissue-specific knockouts were generated by crossing a UAS-Cas9 line with a line expressing 5 UAS-sgRNAs targeting *mor* gene. The specific sequences are provided in the Figure S1A. These flies were then crossed to a tissue-specific GAL4 driver line to induce conditional, clock-neuron-specific knockouts.

### Locomotor activity assays

Locomotor activity of individual adult male flies (3-5 days old) was monitored using the TriKinetics DAM2 Drosophila Activity Monitor system. Flies were placed in glass tubes with 2% agar and 4% sucrose food. Following 5-7 days of LD entrainment, flies were released into DD for 6-7 days. Activity was recorded in 30-minute bins. Rhythmicity, period, and power were analyzed using ClockLab software (Actimetrics) based on the chi-square periodogram (confidence level 0.001).

### In silico protein disorder prediction

The amino acid sequence of *Drosophila* Moira (MOR) was analyzed for intrinsic disorder using the IUPred2A web server (https://iupred2a.elte.hu/). The “long disorder” prediction mode was employed to identify potential intrinsically disordered regions (IDRs) within the full-length protein. This algorithm estimates the capacity of polypeptides to form stable structures based on their total estimated inter-residue interaction energy. A threshold score of 0.5 was used to define disorder; residues with a score exceeding 0.5 were classified as having a high propensity for intrinsic disorder. These predicted disordered regions were further correlated with the structural domains of MOR (Chromo, SWIRM, and SNAT) to define the boundaries of IDR1 (residues 265–424), IDR2 (residues 773–919), and IDR3 (residues 1040–1189) for subsequent in vitro assays.

### In vitro protein expression, purification, and phase separation assay

Molecular cloning: Full-length MOR and its three individual Intrinsically Disordered Regions (IDRs) were cloned into pET28a vectors using Gibson Isothermal Assembly (NEBuilder HiFi DNA Assembly Master Mix). This vector adds an N-terminal 6xHis-SUMO tag. All plasmids were verified by Sanger sequencing.

#### Protein expression and purification

Plasmids were transformed into E. coli BL21 competent cells26262626. A single colony was cultured overnight, reinoculated, and grown to log phase before protein expression was induced with 1 mM IPTG at 16°C for 16 hours. Bacterial pellets were resuspended in high-salt lysis buffer (500 mM NaCl, 50 mM Tris pH 8.0, 10 mM MgCl2, 1% Triton X-100, 25 mM Imidazole, 1 mM DTT, 0.1 mg/mL lysozyme, 1x Benzonase nuclease). The supernatant, collected after centrifugation (20,000 x g, 15 min, 4°C), was incubated with Ni-NTA affinity resin (HisPur Ni-NTA Spin Columns). After 3 washes, the 6xHis-SUMO-tagged MOR protein was eluted with an elution buffer containing high imidazole. Imidazole was removed using Vivaspin concentration tubes (10 kDa MWCO). The 6xHis-SUMO tag was cleaved by incubation with TEV protease overnight at 4°C. The reaction mix was passed through Ni-NTA resin again to remove the tag and TEV enzyme, and the flow-through containing the purified MOR protein was collected. Protein was concentrated, mixed with 10% glycerol, snap-frozen, and stored at -80°C.

#### In vitro LLPS assay

Purified protein stored in a high-salt buffer (500 mM NaCl) was diluted with a no-salt buffer to the desired final protein and NaCl concentrations to initiate the LLPS reaction.

The mix was immediately added to a coverslip. Droplet formation was observed at room temperature using a confocal microscope.

### Live-cell imaging and FRAP

For live imaging of clock neurons, 5-7 day old flies were used. Brains were dissected in PBS medium under low light and mounted on slides using a double-sided tape spacer. Brains were overlaid with Prolong Glass Antifade mounting medium (ThermoFisher Scientific).

Imaging was performed using a Zeiss LSM800 confocal microscope equipped with an AiryScan super-resolution module (125 nm lateral, 350 nm axial resolution) and a 63x Plan-Apochromat Oil (N.A. 1.4) objective. Z-stack series (approx. 250 nm per slice) were collected.

For Fluorescence Recovery After Photobleaching (FRAP) analysis, a defined region of interest (ROI) within an EGFP-MOR focus was photobleached using a high-intensity laser pulse. Fluorescence recovery within the ROI was monitored by time-lapse imaging every few seconds for several minutes. Mean Square Displacement (MSD) analysis was performed by tracking the 3D position of individual foci over time using FIJI.

### DNA fluorescence in situ hybridization (DNA-FISH)

DNA-FISH was performed similarly to previously described protocol^9^. Probes targeting specific genomic loci (*per*, *pdm3*, *Sdc*, *CG10384*, *A2bp1*, 14-3-3ζ, *Argk1*) were designed. Adult brains were dissected, fixed, and permeabilized. Samples were subjected to harsh denaturation steps using formamide at high temperatures before probe hybridization. Following hybridization and washes, samples were mounted and imaged on the Zeiss LSM800 AiryScan system.

### Hybridization chain reaction (HCR) RNA-FISH

HCR-FISH was performed as previously described^47, 48^. Brains were dissected in PBS, fixed for 20 min in 4% PFA, washed, and permeabilized overnight in 70% ethanol at 4°C. Brains were then hybridized overnight at 37°C with probe sets (Molecular Instruments) targeting introns (for nascent transcription of *tim*, *pdm3*, *shaker*) or exons (for mature mRNA of *tim*, *pdp1*). Following washes, brains underwent overnight amplification at room temperature using fluorescently labeled amplifier hairpins. Samples were imaged on the Zeiss LSM800 AiryScan system.

### Combined protein and HCR-FISH imaging

To visualize fluorescently-tagged proteins (e.g., EGFP-MOR) and nascent transcripts simultaneously, we used a modified HCR-FISH protocol designed to preserve fluorescent protein signals. Briefly, brains were dissected and incubated in Prolong Glass Antifade mounting medium for 10 minutes at room temperature before PFA fixation. The standard HCR-FISH protocol (described above) was then followed. This allowed for the co-visualization of EGFP-MOR foci and intron-specific HCR-FISH signals for *tim* and *pdm3*.

### Clock neuron isolation by FACS

For ATAC-seq and RNA-seq, clock neuron nuclei were isolated based on the protocol from a published report^6^. *Clk-GAL4*; *UAS-NLS-GFP* flies were used. Approximately 60 brains were dissected per sample. Brains were dissociated using collagenase and trituration. DAPI was used as a dead cell marker. Live, GFP-positive nuclei (DAPI-negative, GFP-positive) were sorted using a BD Aria III FACS sorter. GFP-negative cells were also collected as controls. Typically, 1,500-3,000 clock neurons were obtained per sample.

### ATAC-seq and RNA-seq analysis

ATAC-seq was performed on FACS-sorted clock nuclei following the protocol from our previous article^6^. Nuclei were pelleted and resuspended in a transposition solution containing Tn5 transposase (Diagenode). Following transposition, DNA was purified. Libraries were prepared using limited-cycle PCR with indexed primers. Libraries were sequenced on a NovaSeq S4 flow cell. Reads were trimmed and aligned to the dm6 genome using Bowtie2. Duplicates and mitochondrial reads were removed. Peak calling was performed using MACS2. Differential accessibility analysis between time points (CT24 vs. CT36) or genotypes (mor-RNAi vs. WT) was performed using DESeq2. GO analysis was performed using FlyEnrichr. RNA was extracted from FACS-sorted clock neurons collected at CT24, CT30, CT36, and CT42. Libraries were prepared and sequenced. Analysis of oscillating transcripts was performed using the RAIN algorithm. GO analysis was performed using FlyEnrichr.

### Quantification and statistical analysis

All image analysis was performed using FIJI(ImageJ) or Zeiss ZEN software.

#### Foci Analysis

Foci intensity, number, and distance to the nuclear envelope were quantified as described in a previous report^9^. Briefly, ROIs were manually drawn around foci or the nucleus. Foci intensity was calculated as the integrated intensity after background subtraction. Distance from the foci centroid to the nearest point on the nuclear envelope polygon was calculated in 3D.

FISH Analysis: HCR-FISH spot intensity was measured as integrated intensity. DNA-FISH gene locus distance to the nuclear envelope was measured similarly to foci distance.

#### Statistical Analysis

Statistical significance was determined using a two-sided Student’s t-test followed by post-hoc analysis, as appropriate. Significance levels are denoted in figures as *p < 0.05, **p < 0.01, ***p < 0.001, and ****p < 0.0001. Data are presented as mean ± SEM.

### RNA extraction and RT-qPCR

To analyze mor mRNA levels, total RNA was extracted from fly heads collected at different circadian time points (CT) in constant darkness. ∼30-50 heads were used per sample. RNA extraction was performed using Trizol reagent (Invitrogen) according to the manufacturer’s protocol. cDNA was synthesized from 1 µg of total RNA using a SuperScript3 reverse transcription kit. Quantitative real-time PCR (qPCR) was performed using a SYBR Green master mix. Relative mRNA levels were calculated using the delta 2 method, normalizing to a housekeeping gene rp49.

## DATA REPOSITORY

RNA-seq and ATAC-seq data are deposited in GSE300582 and GSE300583 repositories.

## ACKNOWLEDGEMENTS

We thank members of the Yadlapalli laboratory for helpful discussions and Yangbo Xiao for help with the design of Ote-GFP CRISPR-edited flies and Christopher Wilson for help with initial set of experiments. We thank the Bloomington Drosophila Stock Center for providing fly strains. This work was supported by National Institutes of Health NIGMS R35GM133737 grant and NINDS RO1NS140761 grant to S.Y., the Alfred P. Sloan fellowship, the McKnight scholar grant, and the Chan Zuckerberg collaborative pairs grant to S.Y., the Rackham International Student Fellowship to Q.Q., and the Rackham Predoctoral Fellowship to Y.Y..

## AUTHOR CONTRIBUTIONS

Q.C. performed all the imaging experiments and RNA-seq experiments and analyzed the data. Y.Y. performed the ATAC-seq experiments. D.C. performed live imaging experiments. Q.C. and S.Y. designed the project and experimental plan. Q.C. and S.Y wrote the manuscript with input from all authors.

---

**Figure S1.**
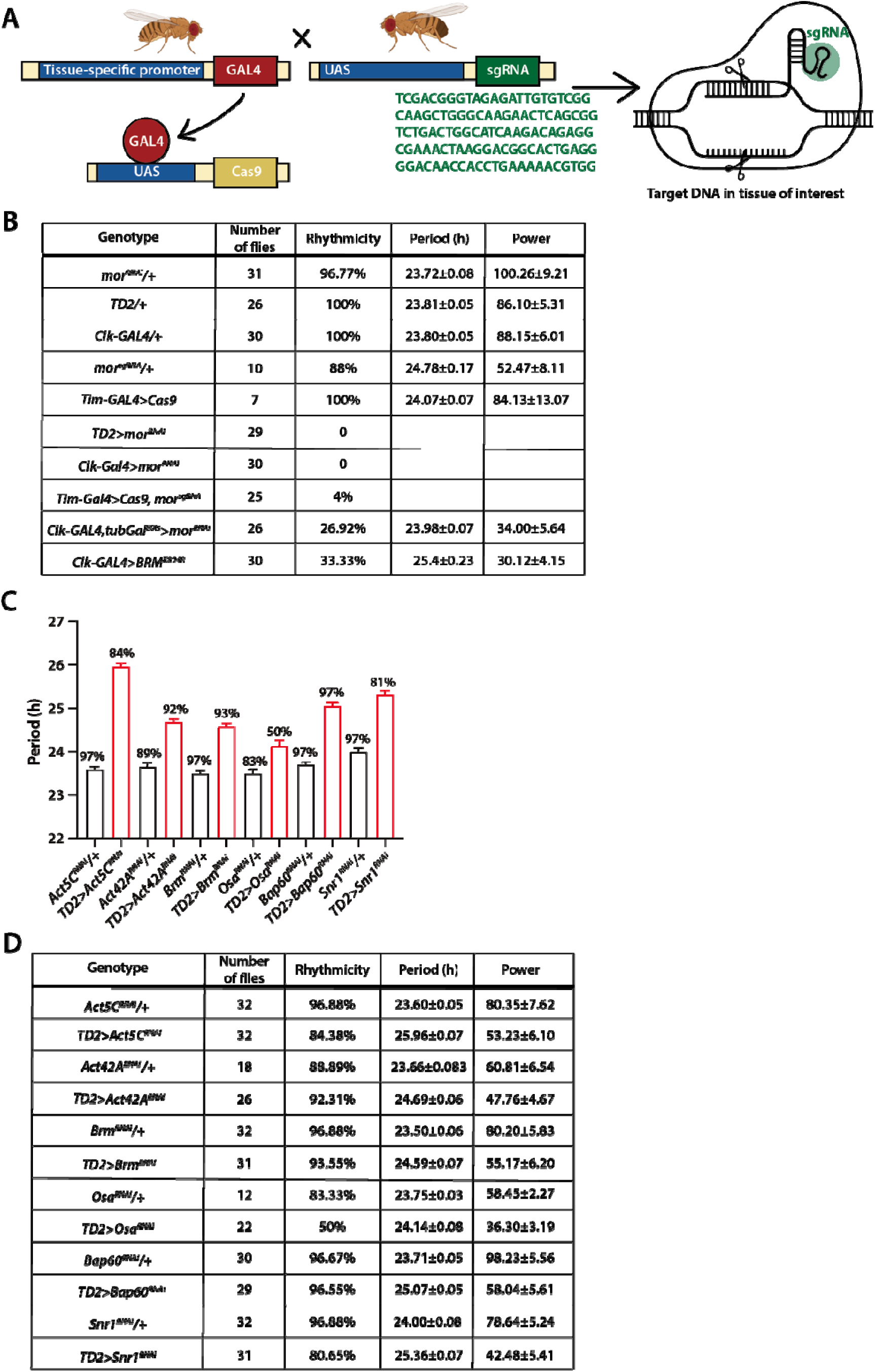
Systematic screen of SWI/SNF subunits identifies MOR as critical for circadian behavior. (A) Schematic diagram of the CRISPR-Cas9 strategy used for generating tissue-specific gene knockouts. A GAL4 driver line is crossed with a *UAS-Cas9* line and a *UAS-mor-sgRNA* line to induce MOR knockout in specific tissues (e.g., clock neurons). sgRNAs used are listed here. (B) Table quantifying locomotor activity for mor perturbations, including mor RNAi driven by multiple GAL4 drivers (severe arrhythmicity, 0% or very low rhythmicity; Fig. 1), adult specific knockdown using *tubGAL^80ts^*, and a dominant negative MOR condition showing partial arrhythmicity. (C-D) Quantification for the knockdown of other core SWI/SNF subunits (Act5C, Act42A, Brm, Osa, Bap60, Snr1) using the *Tim-GAL4, UAS-Dicer2*. These knockdowns show mild to moderate defects, reinforcing that MOR is uniquely critical for robust rhythms.

**Figure S2.**
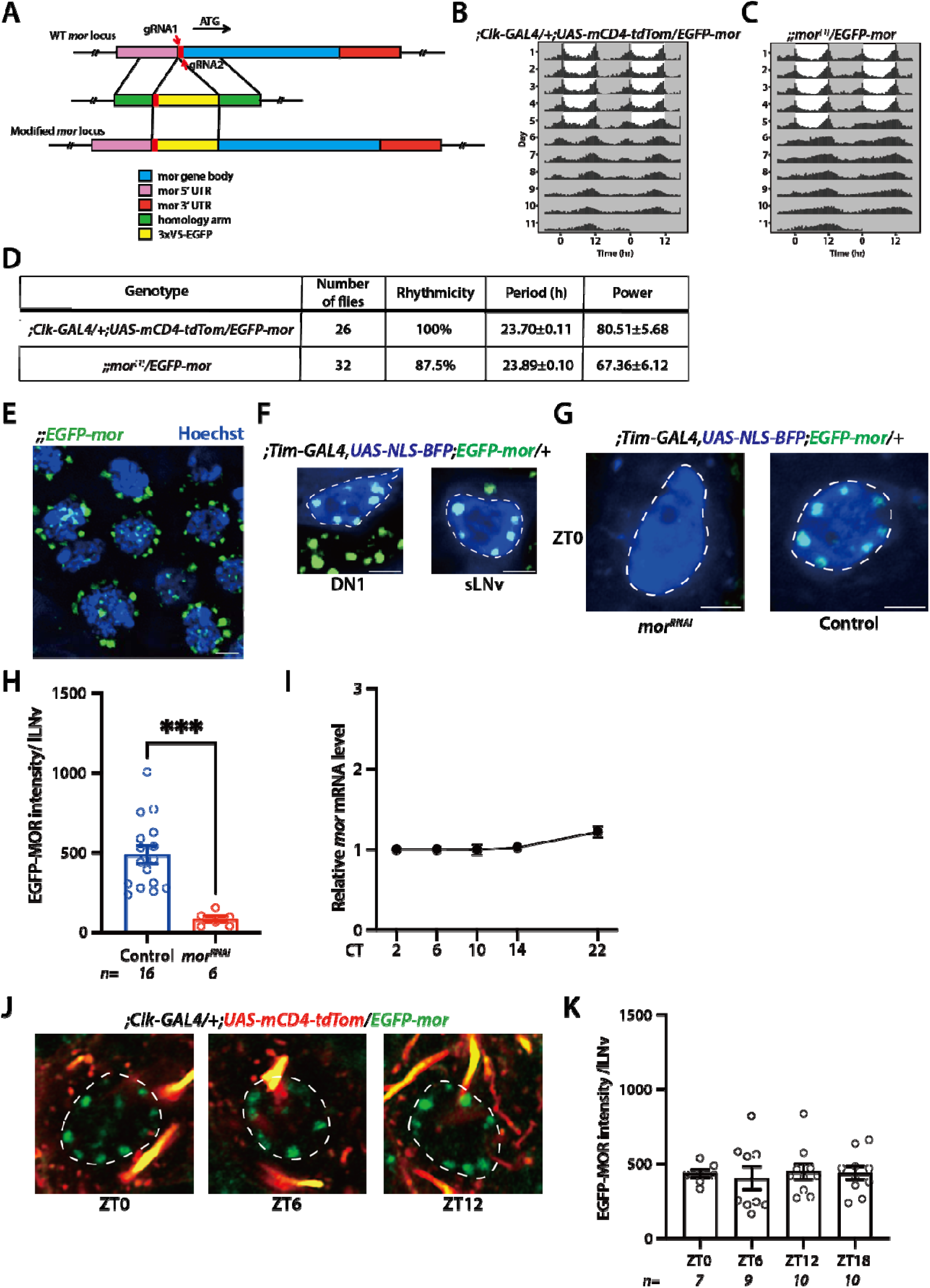
Validation of the functional EGFP-MOR knock-in line, confirmation of mor RNAi efficiency, and non-rhythmic MOR expression in temperature cycle. (A) Schematic of the CRISPR/Cas9-mediated homologous recombination strategy used to tag the endogenous mor locus at its N-terminus with an 3xV5-EGFP tag. (B-D) (B) Locomotor actograms showing that EGFP-MOR flies exhibit fully normal, wild-type rhythmic behavior(left). (C) Flies in which the EGFP-MOR allele is placed over a mor null allele (right) also exhibit robust rhythms, demonstrating that the EGFP-MOR fusion protein functionally rescues the mor mutant phenotype. (E) Representative image of EGFP-MOR foci (green) in multiple nuclei of other cell types (stained with Hoechst, blue). (F) EGFP-MOR foci are present in DN1 (left) and sLNv (right) clock neuron subtypes (nuclei marked by *Tim-GAL4>NLS-BFP*, blue). (G) Representative images of EGFP-MOR (green) in control and *mor^RNAi^* in lLNv neurons. Foci are absent in knockdown condition. (H) Quantification of (G), confirming the high knockdown efficiency of the *mor^RNAi^* line, which results in a significant reduction (***p; 0.001) in EGFP-MOR fluorescence intensity. (I) RT-qPCR analysis of *mor* mRNA levels in fly heads collected over a circadian cycle inconstant darkness (DD). *mor* mRNA is constitutively and non-rhythmically expressed. (J) Representative images of EGFP-MOR foci (green) in lLNv neurons at four different ZTtimes (ZT0, ZT6, ZT12) in temperature cycle (Temperature cycle: 12hr in 18C, 12hr in 25C, with lights off). (K) Quantification of (I), EGFP-MOR intensity in lLNv neurons shows no significant oscillation over a 24-hr temperature cycle.

**Figure S3.**
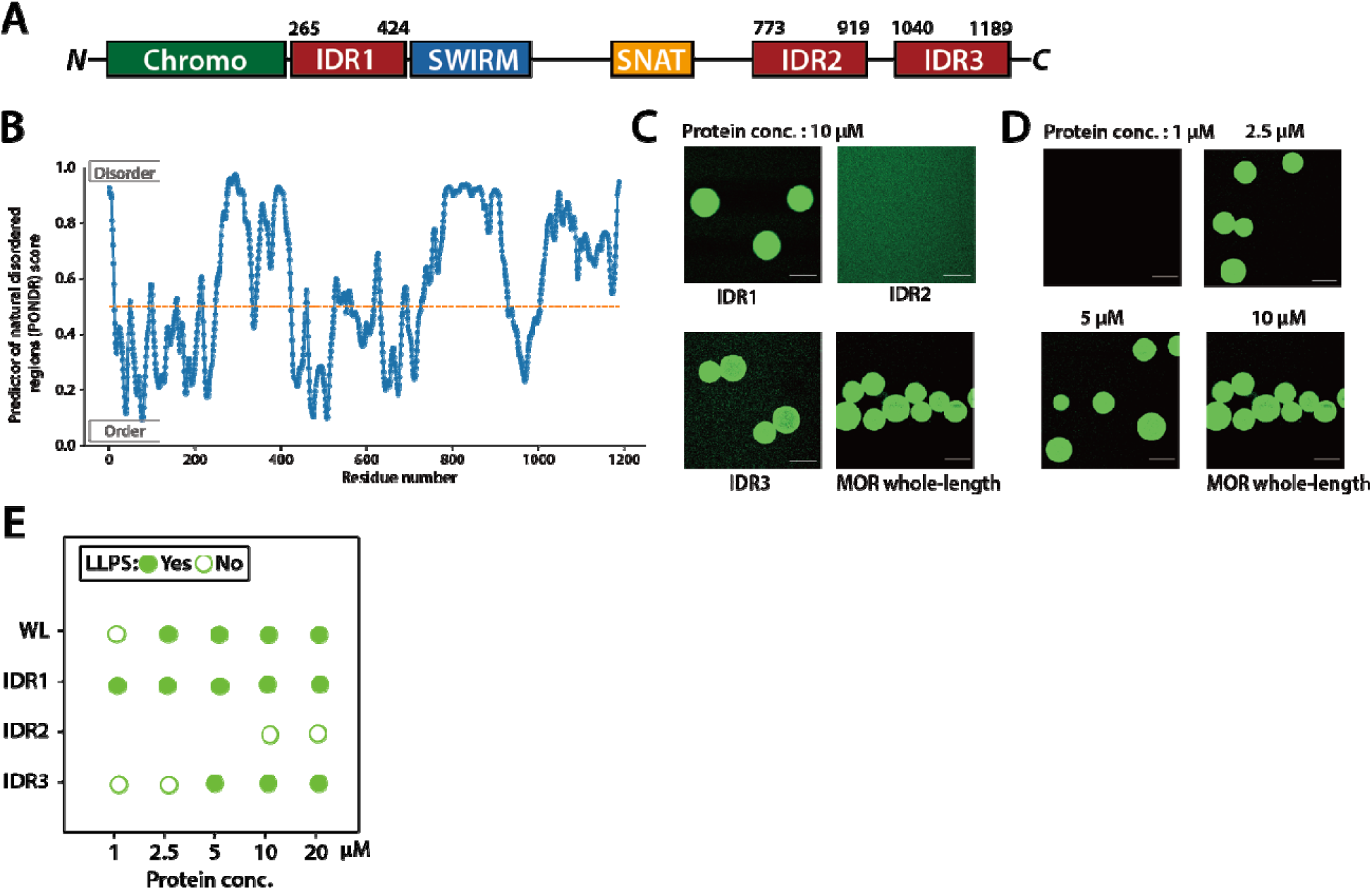
MOR contains Intrinsically Disordered Regions (IDRs) that drive in vitro phase separation. (A) Schematic diagram of the MOR protein domain structure, highlighting three predicted IDRs (IDR1, IDR2, IDR3) in addition to the Chromo, SWIRM, and SNAT domains. (B) Intrinsic disorder prediction for the full-length MOR protein sequence using IUPred2A. Regions with scores above the dashed line (0.5) have a high propensity for disorder, corresponding to the locations of the IDRs. (C) In vitro phase separation assays of purified, fluorescently labeled full-length MOR and its individual IDR fragments. At 10 µM protein concentration, full-length MOR, IDR1, and IDR3 readily form micrometer-scale droplets, while IDR2 remains diffuse. (D) Droplet formation of purified full-length MOR is concentration-dependent, with liquid-liquid phase separation (LLPS) evident at 2.5 µM protein concentration and above. (E) Summary chart of the LLPS properties for whole-length (WL) MOR and its IDRs across a range of protein concentrations. Green circles indicate observed LLPS (droplet formation), and white circles indicate no LLPS.

**Figure S4.**
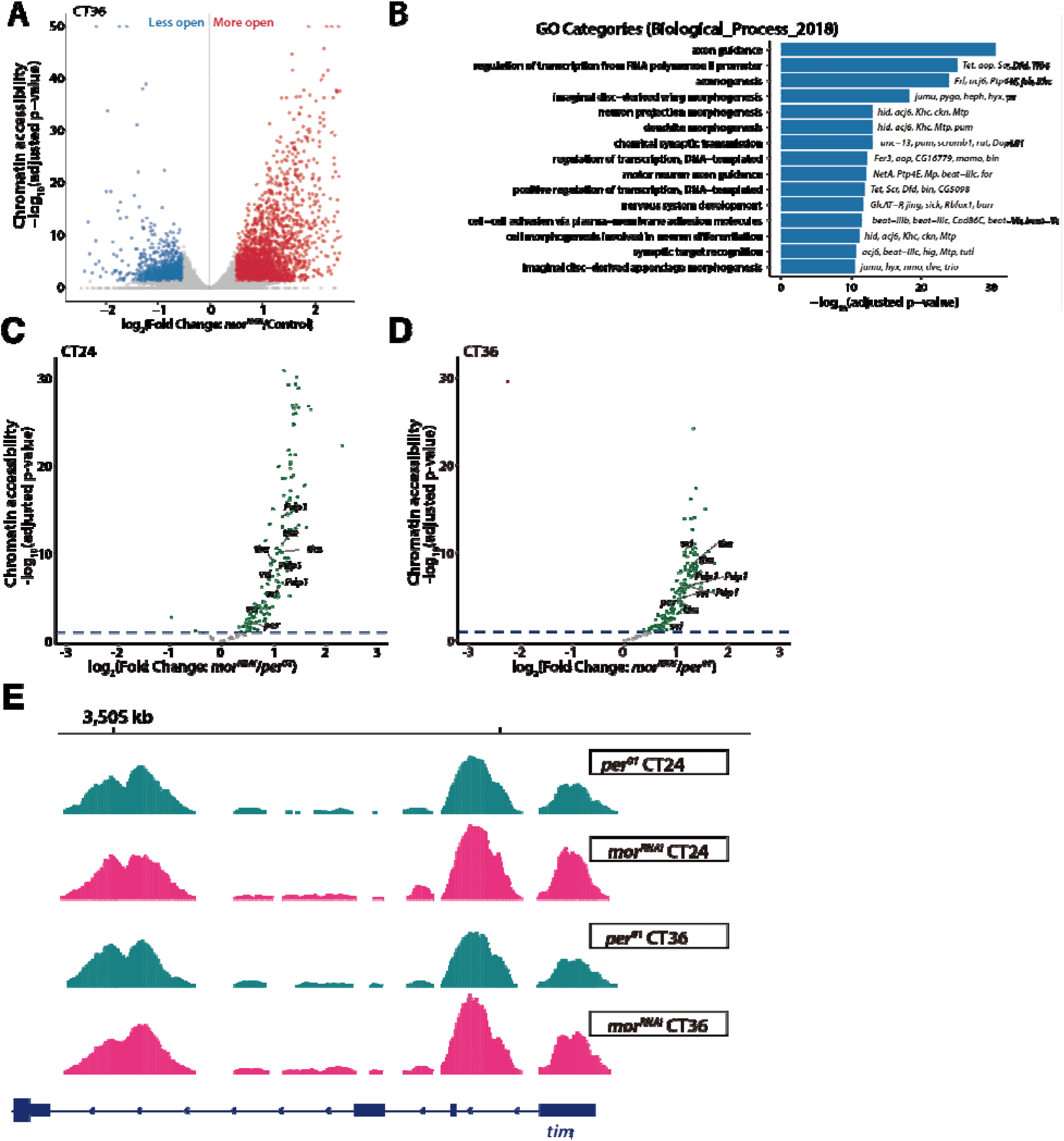
MOR provides a baseline repressive function on chromatin accessibility that is independent of, and stronger than, PER-mediated repression. (A) Volcano plot showing differentially accessible regions (DARs) in *mor^RNAi^* clock neuronscompared to control at CT36. Genes with significantly increased accessibility (more open) are in red, and those with decreased accessibility (less open) are in blue. (B) Gene Ontology (GO) enrichment analysis (2018 Biological Process) for genes associated with the DARs identified at CT36.The most enriched categories relate to transcriptional regulation and nervous system development. (C-D) Volcano plots showing chromatin accessibility changes in *mor^RNAi^* neurons relative to *per^01^* at CT24 (C) and CT36 (D). Green dots indicate genes that peak at CT36 in control (same as Fig. 2E-F). CT36-peaking genes still display elevated accessibility in *mor^RNAi^* flies (log₂FoldChange > 0) compared to *per^01^* both in CT36 and CT24. (E) Representative ATAC signal pile-up tracks at *tim* loci. Signal is normalized to total number of reads for each sample and therefore is comparable among samples. Four different replicates are overlaid. *mor^RNAi^* flies show stronger ATAC signal in *timeless* (*tim)* loci compared to the *per^0^* both in CT36 and CT24.

**Figure S5.**
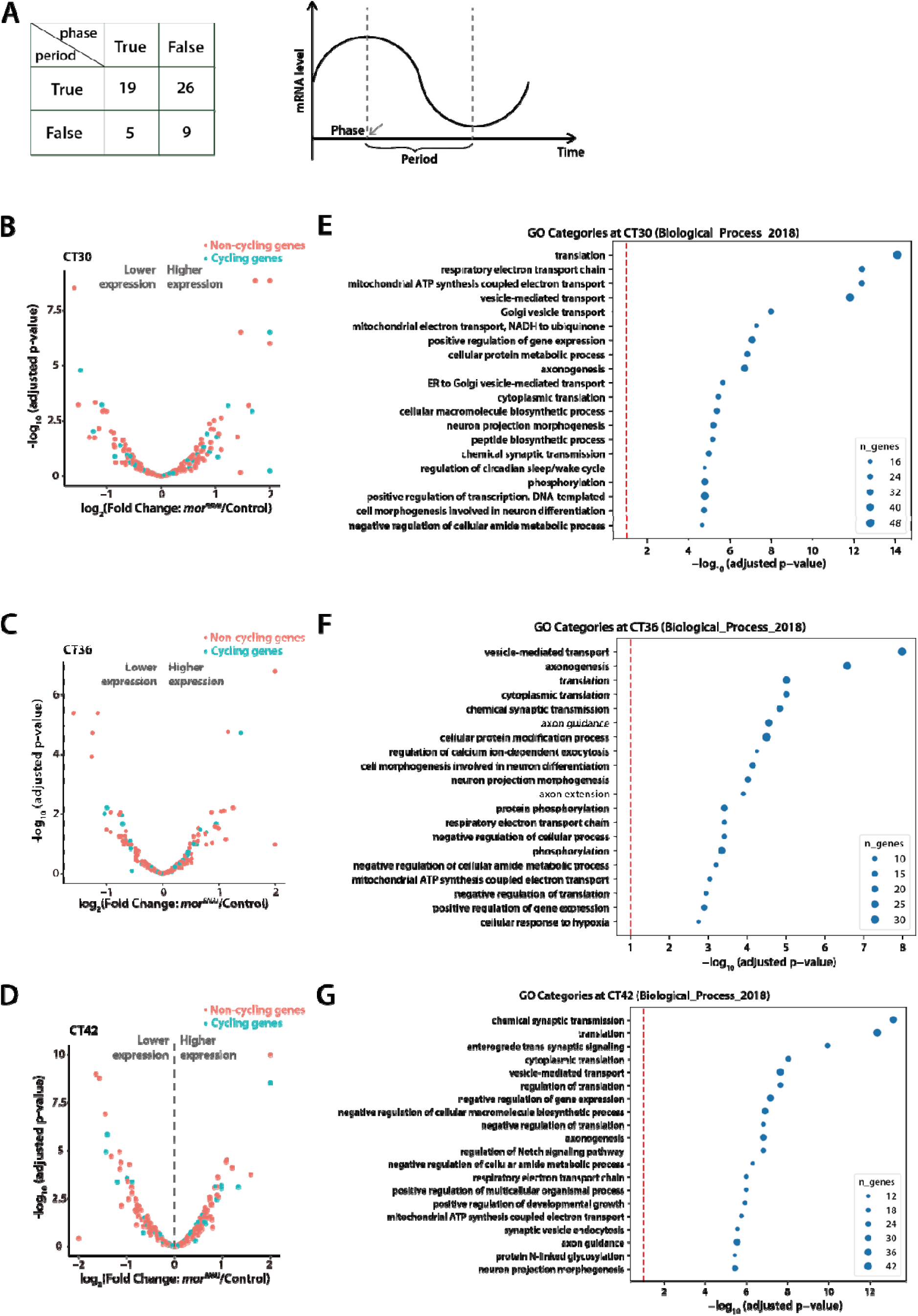
Widespread transcriptional dysregulation in *mor^RNAi^* clock neurons occurs throughout the circadian cycle. (A) A total of 45 transcripts were identified both rhythmic in control and *mor^RNAi^*. However, while they stayed rhythmic in *mor^RNAi^* background, these genes exhibited significant alterations in their phase (time point when mRNA reaches peak expression) and period (period between peak and trough) compared to control. (B-D) Volcano plots showing differential gene expression (*mor^RNAi^* vs. control) at threecircadian timepoints: (B) CT30, (C) CT36, and (D) CT42. These plots complement the CT24 data in Figure 4C. Significant expression changes are evident in both gene populations at all time points. (E-G) Gene Ontology (GO) enrichment analysis (2018 Biological Process) for genes thatwere differentially expressed in mor-RNAi neurons at (E) CT30, (F) CT36, and (G) CT42. Enriched categories consistently include translation, cellular metabolism, and various aspects of neural development and function, indicating broad transcriptional consequences of MOR loss.

**Figure S6.**
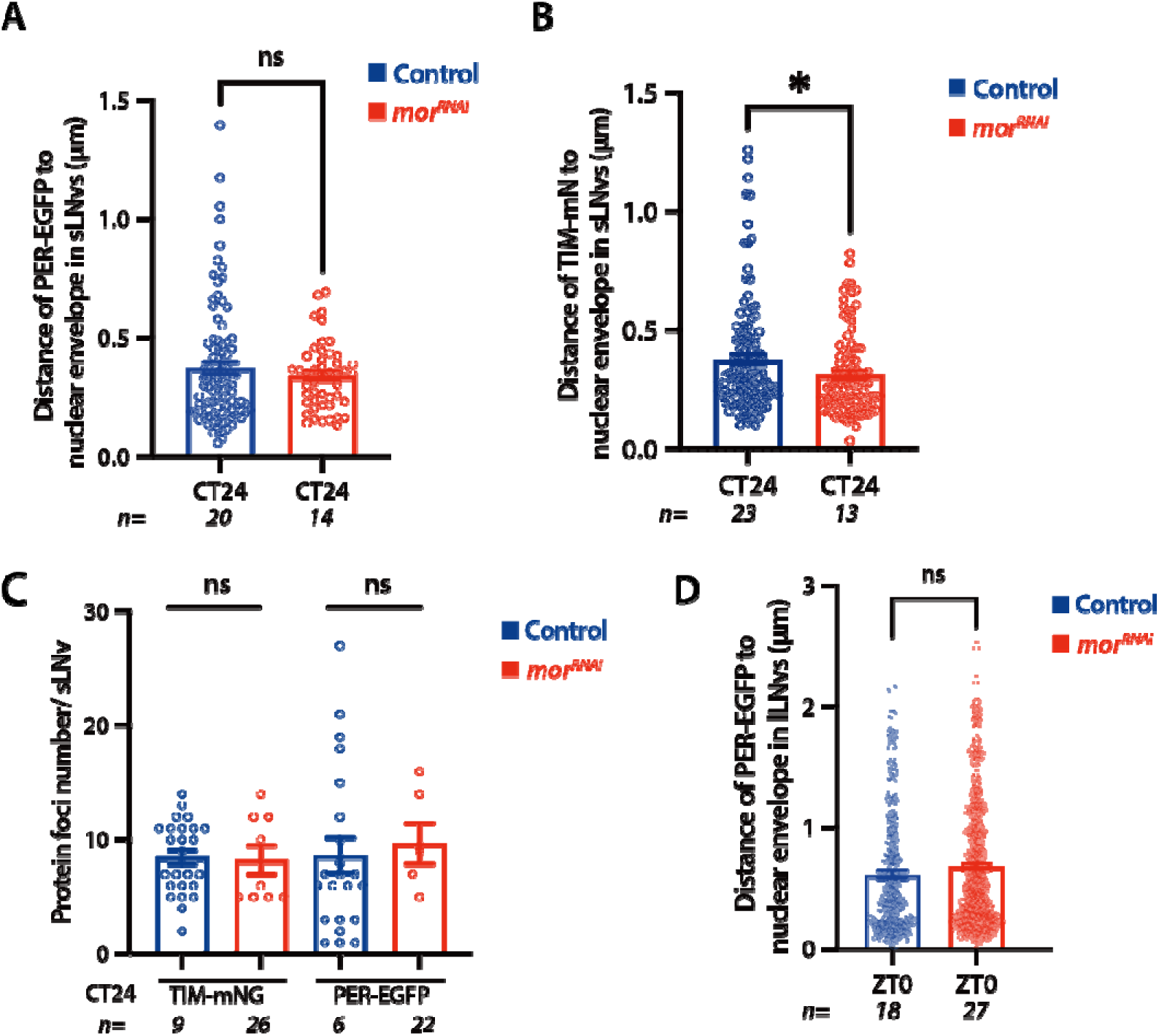
MOR knockdown does not alter the subnuclear localization of PER and TIM foci or their number. (A) Quantification of the distance from PER-EGFP foci to the nuclear envelope in sLNvs at CT24. No significant difference was observed between control (blue) and *mor^RNAi^* (red) flies. (B) Quantification of the distance from TIM-mNG foci to the nuclear envelope in sLNvs at CT24. TIM-mNG foci are positioned slightly closer to the envelope in *mor^RNAi^*flies (*p < 0.05). (C) Quantification of the number of TIM-mNG and PER-EGFP protein foci per lLNv nucleus at CT24. MOR knockdown does not affect the number of foci formed by either protein. (D) Quantification of the distance from PER-EGFP foci to the nuclear envelope in lLNvs at ZT0. No significant difference was observed between control (blue) and *mor^RNAi^*(red) flies. Statistics: Error bars represent mean ± SEM. *p < 0.05, ***p < 0.0001, determined by t-test.

**Figure S7.**
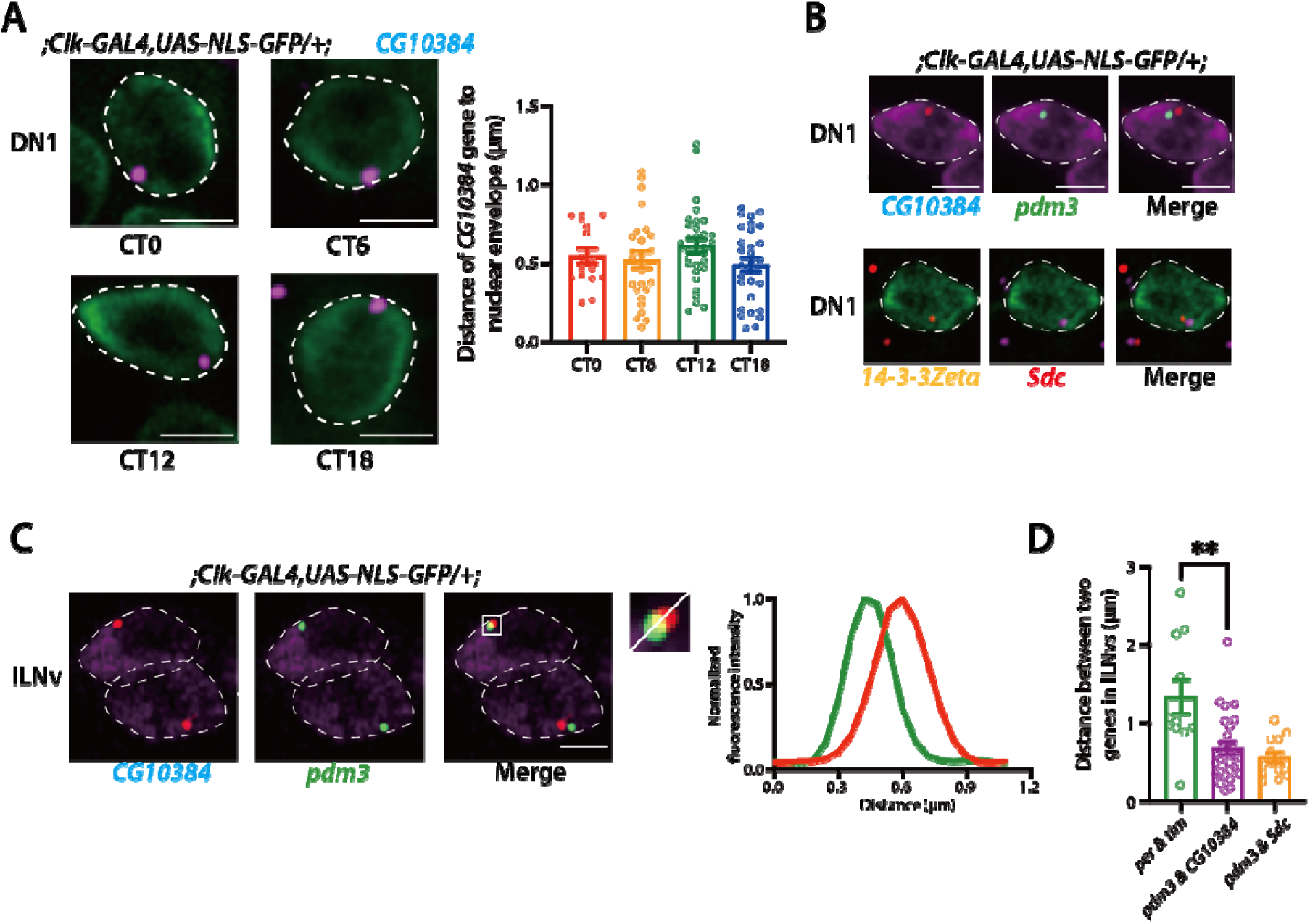
Clock-regulated gene loci show stable peripheral anchoring and chromosome-dependent spatial organization. (A) Representative DNA-FISH images of the *CG10384* locus (magenta) in clock neurons (nucleus labeled by NLS-GFP, green) at four circadian time points (CT0, CT6, CT12, CT18). The graph (right) quantifies the distance of the *CG10384* gene to the nuclear envelope, showing no significant oscillation and confirming its stable peripheral localization. (B) Representative DNA-FISH images showing the clustering of intra-chromosomal gene pairs. (C) Representative DNA-FISH images and corresponding line-intensity distribution plot showing the spatial clustering of the *CG10384* (red) and *pdm3* (green) loci in lLNv. (D) Quantification of the 3D distance between gene pairs in lLNv. The inter-chromosomal pairs *per & tim* are spatially segregated (large distance). The *pdm3 & Sdc* and *pdm3 & CG10384* are pair significantly more clustered in comparison (**p < 0.01). Statistics: Error bars represent mean ± SEM. *p < 0.05, ***p < 0.0001, determined by t-test.

**Figure S8.**
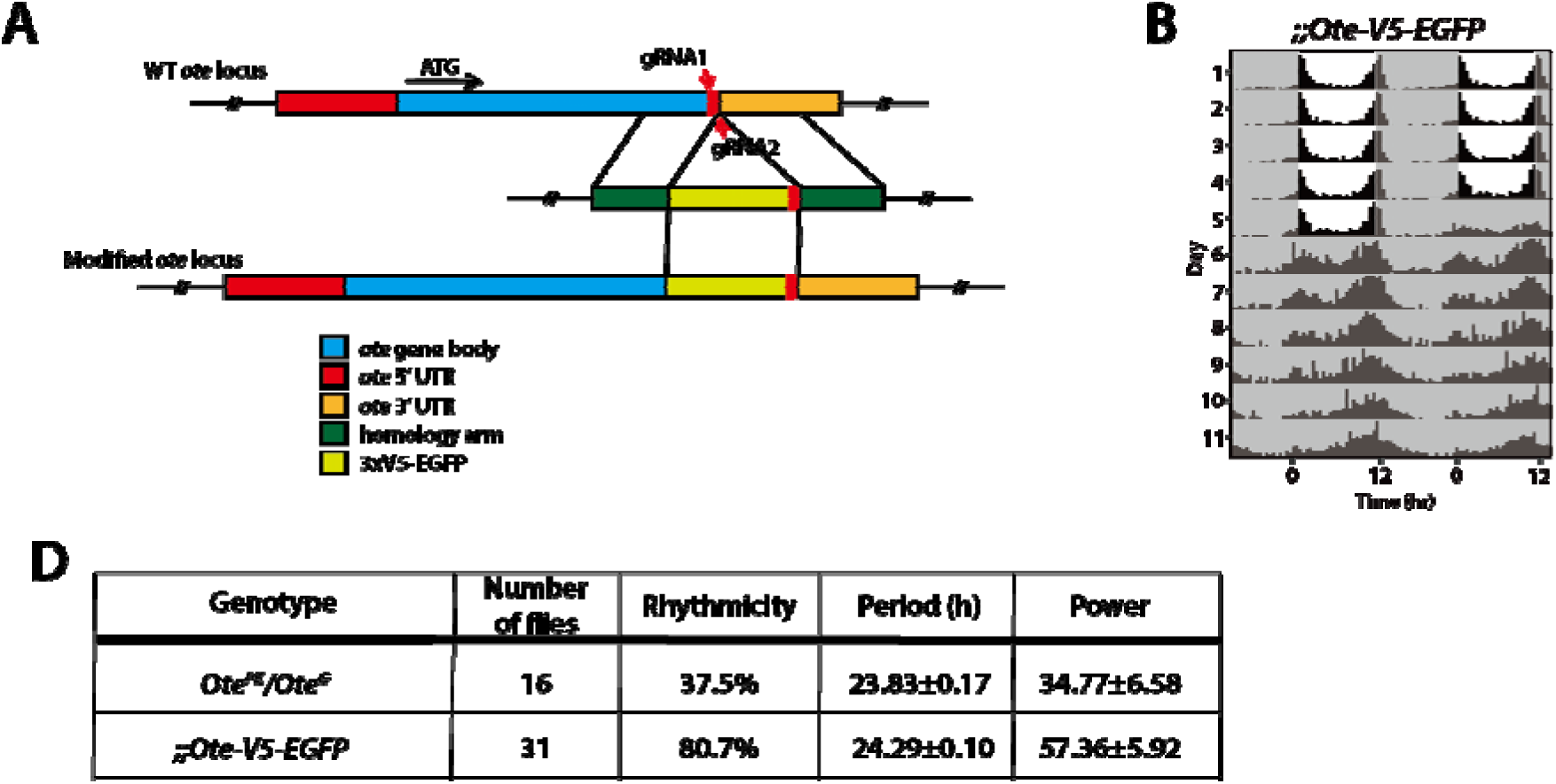
CRISPR/Cas9 generated endogenous tagging of the *otefin* locus. (A) Schematic of the CRISPR/Cas9-mediated homologous recombination strategy used to tag the endogenous *ote* locus at its C-terminus with an 3xV5-EGFP tag. (B) Locomotor actograms showing that *Ote-EGFP* flies exhibit rhythmic circadian behavior. (C) Table quantifying locomotor activity for *Ote^PK^/Ote^G^* (Ote null mutant), Ote-EGFP flies.

